# Three-axis classification of mouse lung mesenchymal cells reveals two populations of myofibroblasts

**DOI:** 10.1101/2021.08.03.454930

**Authors:** Odemaris Narvaez del Pilar, Jichao Chen

## Abstract

The mesenchyme consists of heterogeneous cell populations that support neighboring structures and are integral to intercellular signaling. Despite such importance, mesenchymal cell types are poorly defined morphologically and molecularly, lagging behind their counterparts in the epithelial, endothelial, and immune lineages. Leveraging single-cell RNA-seq, three-dimensional imaging, and lineage tracing, we classify the mouse lung mesenchyme into three proximal-distal axes that are associated with the endothelium, epithelium, and interstitium, respectively. From proximal to distal, (1) the vascular axis includes vascular smooth muscle cells and pericytes that transition as arterioles and venules ramify into capillaries; (2) the epithelial axis includes airway smooth muscle cells and two populations of myofibroblasts: ductal myofibroblasts, surrounding alveolar ducts and marked by CDH4, HHIP, and Lgr6, which persist post-alveologenesis, and alveolar myofibroblasts, surrounding alveoli and marked by high expression of PDGFRA, which undergo developmental apoptosis; (3) the interstitial axis, residing between the epithelial and vascular trees and sharing a newly-identified marker MEOX2, includes fibroblasts in the bronchovascular bundle and the alveolar interstitium that are marked by IL33/DNER/PI16 and Wnt2, respectively. Single-cell imaging reveals distinct morphology of each mesenchymal cell population. This classification provides a conceptual and experimental framework applicable to other organs.

## INTRODUCTION

Cell types are often named after their characteristic features, as in pyramidal neurons and rod/cone photoreceptors for their shape, basal cells and endothelial cells for their location, and cardiomyocytes and hepatocytes for their tissue origin. In contrast, mesenchymal cells, a heterogeneous group of space-filling fibroblasts present in most tissues, lack distinguishing features and remain poorly defined. Even their major embryonic source, the mesoderm, is simply named for its location between the ectoderm and endoderm. Such secondary nature of mesenchymal cells likely reflects the fact that they often do not form a recognizable structure on their own and instead support other cell lineages, such as epithelial and endothelial tubes. This supportive role suggests to us that mesenchymal cells may be defined relative to their better understood neighbors – a concept applied in this study.

The typical challenge in defining mesenchymal cells despite their importance is exemplified in the mammalian lung. The lung mesenchyme provides chemical signals and physical constraint necessary for branching and alveolar morphogenesis, forms niches for airway and alveolar stem cells, and goes awry during fibrosis and tumorigenesis (Lambrechts et al., 2018; McCulley et al., 2015; Zepp et al., 2017). However, unlike the epithelial, endothelial, and immune lineages where major cell types have been largely defined (Tabula Muris et al., 2018; Travaglini et al., 2020), a consensus of the diverse mesenchymal cell types has yet to emerge. As the aforementioned cell-type-defining features are less forthcoming for mesenchymal cells, the field has relied on components of signaling pathways of known importance in mesenchymal biology, such as Pdgf (Endale et al., 2017; Li et al., 2018; Muhl et al., 2020), Fgf (El Agha et al., 2014; Hagan et al., 2020), Shh (Cassandras et al., 2020; Kugler et al., 2017; Li et al., 2015), and Wnt (Lee et al., 2017; Zepp et al., 2017) pathways. Due to the dynamic nature of signaling pathways and their deployment in multiple concurrent processes, mesenchymal cell types tagged by candidate signaling molecules might not align with those classically defined by molecular and cellular criteria.

Recent accumulation of single-cell genomic data has allowed unbiased identification of lung mesenchymal cell types (Liu et al., 2021; Riccetti et al., 2020; Tsukui et al., 2020; Xie et al., 2018). However, computationally distinct cell types need to be mapped to the native tissue to integrate the molecular differences with their anatomical and cellular differences. This mapping is challenging in the lung due to its complex, three-dimensional (3D) structure where (1) topologically distal regions may be in the center of a section and (2) the cell body and its ramifying processes may appear discontinuous on a section. These issues can be alleviated by whole-mount immunostaining and 3D imaging, as we have learned from our recent work on the expansive alveolar type 1 cells and identification of *Vegfa* and NKX2-1 expression in them (Little et al., 2019; Yang et al., 2016). On the other hand, single-cell genomics now allows cells that have been named after their cellular features, such as lipofibroblasts and matrix fibroblasts (McGowan and Torday, 1997; Xie et al., 2018), to be examined for accuracy based on the corresponding lipid and matrix-related molecules.

In this study, we posit that lung mesenchymal cell types can be mapped relative to the epithelial and endothelial trees they support and that antibody-based 3D imaging can distinguish intermingled cells and their processes to identify cell-type-specific morphology. The resulting molecular, anatomical, and cellular features allow us to classify the mouse lung mesenchyme into three axes – vascular, epithelial, and interstitial – that are each partitioned proximal-distally with cells of distinct morphology. This classification reveals two populations of myofibroblasts that respectively surround the alveolar ducts or the alveoli and persist or disappear post-alveologenesis.

## RESULTS

### Time-course scRNA-seq identifies known and new mesenchymal cell populations that organize into vascular, epithelial, and interstitial axes

Without a consensus surface marker to sort lung mesenchymal cells, we used a negative-gating strategy to separate the other cell lineages, namely epithelial, endothelial, and immune lineages that were labeled by CDH1 (also known as E-Cadherin), ICAM2, and CD45 (official name PTPRC), respectively (Fig. S1A). We previously showed that this cell isolation strategy allowed balanced, sufficient sampling of all major cell types in both developing and mature mouse lungs (Cain et al., 2020; Vila Ellis et al., 2020). We collated single-cell RNA-seq (scRNA-seq) data of lungs spanning embryonic, neonatal, juvenile, and adult stages, and readily identified the mesenchymal cells as positive for a matrix gene *Col3a1* and negative for other cell lineage markers including *Nkx2-1* (epithelial), *Cdh5* (endothelial), and *Ptprc* (immune) (Fig. S1B).

We used Harmony (Korsunsky et al., 2019) to align comparable cell types across time points without changing the raw gene expression values that we would use for subsequent differential analysis. To reduce bias, we performed unsupervised clustering and numerically named the resulting 24 cell clusters, and then annotated them based on temporal dynamics and known marker genes (Fig. 1A, B, Table S1). Developmental-stage-specific cell clusters were [1] specific to embryonic day (E) 17 and thus presumably less differentiated progenitors that were clustered with their mature counterparts (clusters 19 and 22); [2] proliferative as it was largely limited to before postnatal day (P) 7 and expressed a proliferation marker *Mki67* (clusters 11, 14, and 21); or [3] developmentally cleared as it disappeared between P13 and P20 (cluster 6 and the majority of cluster 1).

**Figure 1:**
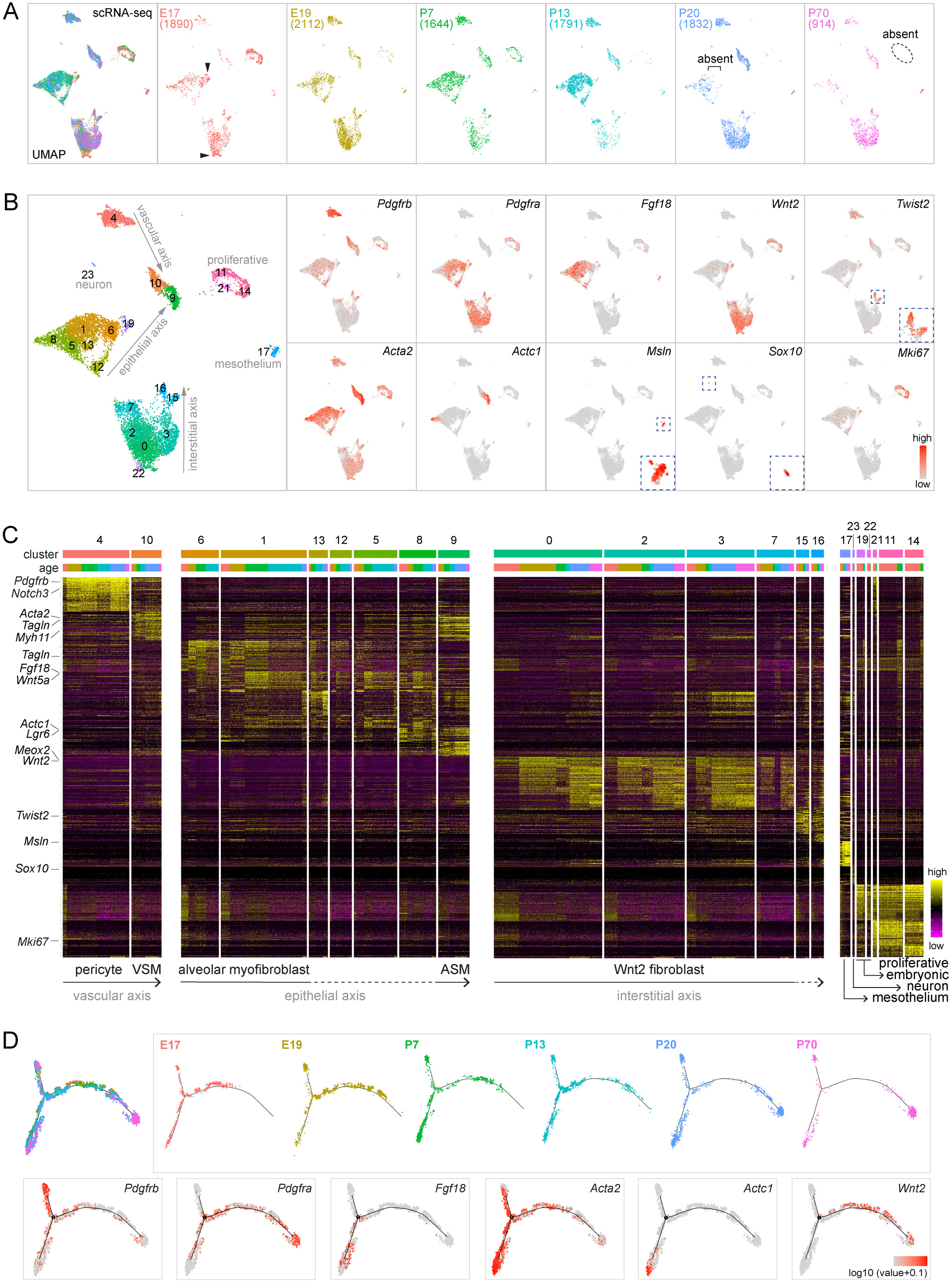
Time-course scRNA-seq identifies known and new lung mesenchymal cell populations that organize into vascular, epithelial, and interstitial axes. (**A**) ScRNA-seq UMAPs of lung mesenchymal cells across developmental ages with cell numbers in parenthesis. Embryonic clusters (arrowhead) are absent postnatally. Proliferative cells (dashed oval) are absent in the adult lung (P70). Cells marked by the square bracket are absent in mature lungs (P20 and P70). (**B**) Seurat unbiased clustering groups lung mesenchymal cells into 24 clusters. Established markers identify pericytes (*Pdgfrb*; cluster 4), vascular smooth muscle cells (*Acta2/Pdgfrb*; cluster 10) airway smooth muscle cells (*Acta2/Actc1*; cluster 9), mesothelial cells (*Msln*; cluster 17), neurons (*Sox10*; cluster 23), and proliferative cells (*Mki67*; clusters 11, 14, 21). *Wnt2* and *Twist2* mark transcriptionally related populations. *Acta2* is high in airway and vascular smooth muscle cells (ASM and VSM for clusters 9 and 10, respectively) and myofibroblasts that are marked by *Pdgfra* and *Fgf18*. Embryonic clusters in (**A**) are numbered 19 and 22 and express *Wnt2*. The proposed 3 axes are labeled. (**C**) Marker heatmaps with clusters in (**B**) grouped by transcriptional similarity and ordered by ages within each cluster. Clusters of unidentified cell types are marked by dashes in the arrow. (**D**) Monocle coalesces lung mesenchymal cells into 3 trajectories whose termini correspond to ASM cells (*Actc1*), pericytes (*Pdgfrb, Acta2-*), and Wnt2 fibroblasts (*Wnt2*), respectively. Cells from mature lungs (P20 and P70) are largely restricted to the termini.

Known mesenchymal cell types were expectedly captured in our dataset, including pericytes marked by *Pdgfrb* (cluster 4), vascular and airway smooth muscle (VSM and ASM) cells marked by *Acta2* and distinguishable by *Pdgfrb* and *Actc1* (Ijpma et al., 2020), respectively (clusters 10 and 9), and myofibroblasts marked by *Acta2, Pdgfra* and *Fgf18* albeit with additional heterogeneity to be addressed later (clusters 1, 5, 6, 8, 12, and 13) (Fig. 1C). The remaining major clusters formed a large group (clusters 0, 2, 3, and 7) and a small group (clusters 15 and 16) marked by *Wnt2* and *Twist2* (also known as *Dermo1*), respectively. These two groups were recently named after additional markers as Col13a1 and Col14a1 matrix fibroblasts (Xie et al., 2018), although most mesenchymal cells as well as endothelial and epithelial cells produced matrices (Fig. S1C). Via the process of elimination and based on their abundance, the *Wnt2* and *Twist2*-expressing cells should include lipofibroblasts that were considered to have unique lipid metabolism (McGowan and Torday, 1997); however, the commonly used marker *Plin2* (also known as *Adrp*) was non-specific, raising questions on the accuracy of the prefix “lipo” (Fig. S1C). We additionally identified the less abundant mesothelial cells (cluster 17; marked by *Msln*) and even rare neurons (cluster 23; marked by *Sox10*).

As reasoned in the introduction, we hypothesized that mesenchymal cells could be classified based on the structures they supported, most notably the vascular and epithelial trees. It was self-evident to assign vascular smooth muscle cells and pericytes to the vascular tree, which we named the vascular axis because the two mesenchymal cell types situated along a proximal-distal axis. Applying the same concept, we assigned to the epithelial axis the proximal ASM cells (cluster 10) and the distal alveolar myofibroblasts (clusters 6 and majority of 1), both of which constrain and shape the epithelium (Kim and Vu, 2006), and predicted that the other transcriptionally-related clusters (5, 8, 12, and 13) were associated with the epithelium in-between, namely the alveolar ducts – for which we provided evidence later in this study. Finally, recognizing that the *Twist2-expressing* cells were shown in a mouse phenotyping database (Koscielny et al., 2014) to localize between the epithelial and endothelial trees within the proximal bronchovascular bundles and that the ratio of *Twist2*-expressing and *Wnt2*-expressing cells was what one would expect for proximal and distal compartments, we predicted that they belonged to a third axis and named it the interstitial axis to refer to the space between the epithelial and endothelial trees. As a result, the vascular, epithelial, and interstitial axes, each arranged proximal-distally, accounted for all the major cell clusters with their corresponding markers (Fig. 1B, C). The proliferative clusters (11, 14, and 21) included cells from each of the three axes as cell cycle genes dominated over cell type markers.

Supporting this three-axis classification, Monocle trajectory analysis coerced the associated mesenchymal cells into three paths that terminated in *Pdgfrb*+ pericytes, *Actc1*+ ASM cells, and *Wnt2*+ fibroblasts – corresponding to the vascular, epithelial, and interstitial axes, respectively (Fig. 1D). Reciprocally, cells from the three Monocle trajectories largely mapped to the proposed three axes on the Uniform Manifold Approximation and Projection (UMAP) (Fig. S2). Although the root of Monocle trajectories was arbitrary (Trapnell et al., 2014), cells from the mature lungs (P20 and P70) were largely limited to the three termini, whereas cells from developing lungs were more centrally located, suggesting gradual specialization along the three axes. Below, we focused on each axis individually to define constituent cell populations, map in 3D their proximal-distal distributions, and categorize cell morphology.

### Within the vascular axis, proximal vascular smooth muscle cells transition to distal pericytes, which mature postnatally

Reclustering of cells of the vascular axis (Fig. 1C) identified one VSM population, marked by contractile genes *Acta2* and *Tagln*, and three populations of pericytes – proliferative, immature, and mature (Fig. 2A, Table S2). While all cells of the vascular axis expressed *Pdgfrb* and *Notch3*, immature pericytes mainly included cells from E17, E19, and P7 lungs and expressed a higher level of ribosomal genes (*Rplp0* and *Rpl7a*) possibly to support cell growth, whereas mature pericytes mostly had cells from P13, P20, and P70 and expressed more *Gap43* and *Gucy1a1* (Fig. 2A) – suggesting that pericytes, unlike VSM cells, mature after birth.

**Figure 2:**
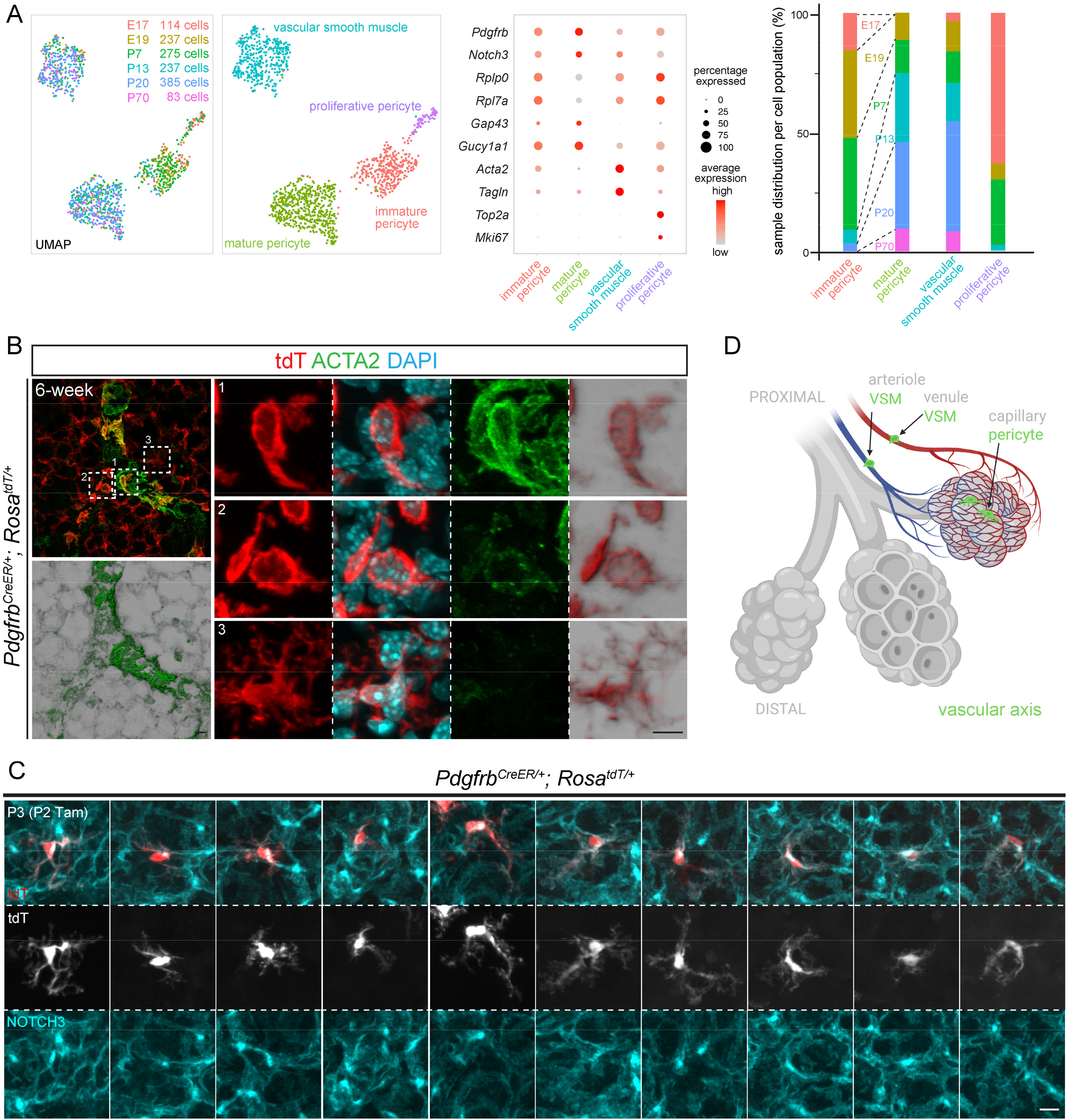
Within the vascular axis, proximal vascular smooth muscle cells transition to distal pericytes, which mature postnatally. (**A**) ScRNA-seq UMAPs of lung mesenchymal cells of the vascular axis, colored for developmental ages (left) or cell types (right), as supported by relevant markers in the dot plot. Immature pericytes are mostly from embryonic (E17 and E19) and neonatal (P7) lungs, as proliferative pericytes are (bar graph). (**B**) Immunostaining images of a transition zone between VSM cells and pericytes that are genetically labeled by *Pdgfrb^CreER^* with 3 mg tamoxifen administered 48 hr before lung harvest. Intermediate cells (box 2) have low ACTA2 but the morphology of VSM cells (box 1) and not that of pericytes (box 3). Scale: 10 um. Images are representative of at least three biological replicates (same for all subsequent images). (**C**) Single cell morphology of 10 pericytes from sparse genetic labeling. Tam, 250 ug tamoxifen. Scale: 10 um. (**D**) Diagram showing VSM cells and pericytes of the vascular axis. Created with BioRender.com.

To visualize the vascular axis, we used a *Pdgfrb^CreER^* driver (Cuervo et al., 2017) and wholemount immunostaining to identify the transition from VSM cells to pericytes based on the tube diameter expected for arterioles and the gradual decrease in ACTA2 (also known as SMA) staining (Fig. 2B). The *Rosa^tdT^* reporter readily defined the nucleus and cell morphology and showed that cells with both high and low ACTA2 expression had nucleus-length, strained cellular projections often at an oblique angle to the vessels, different from the circumferential wrapping of airways by ASM cells (Fig. 2B). In contrast, ACTA2-negative pericytes had long astrocyte-like extensions that were often polarized toward one side of the nucleus and decorated with NOTCH3, which was a pericyte marker and also concentrated in a perinuclear vesicular compartment as PDGFRB was (Huang et al., 2007) – a subcellular distribution to be taken into account when enumerating pericytes (Fig. S3A, 2C). Therefore, the vascular axis consists of proximal VSM cells and distal pericytes sharing PDGFRB/NOTCH3 albeit with contrasting morphology (Fig. 2D).

### The epithelial axis includes airway smooth muscle cells as well as two populations of myofibroblasts, marked by CDH4/HHIP/LGR6, and high PDGFRA, respectively

Next, we subsetted and reclustered cells of the epithelial axis, and readily identified an *Actc1*+ ASM cluster that was largely unchanged over time and an *Acta2/Tagln/Myh11*+ myofibroblast cluster that shifted on the UMAPs, reflecting changes in gene expression and/or cell composition (Fig. 3A, Table S3). Most notably, at P7 and P13, the known PDGFRA-high alveolar myofibroblasts (Endale et al., 2017) only constituted part of the cluster and were adjacent to a second *Cdh4/Hhip/Lgr6*+ population (Fig. 3A). Intriguingly, *Cdh4/Hhip/Lgr6*+ cells remained in the same UMAP location at P20 and P70, when the PDGFRA-high alveolar myofibroblasts disappeared (Fig. 3A). These two populations of myofibroblasts – albeit at different UMAP locations, possibly as immature precursors – were also distinguishable at E17 and E19 based on *Cdh4* and *Lgr6* expression, although *Pdgfra* and *Hhip* expression was non-discriminatory embryonically and only became restricted to their respective myofibroblast populations postnatally (Fig. 3A). Below, we detailed the two myofibroblast populations individually.

**Figure 3:**
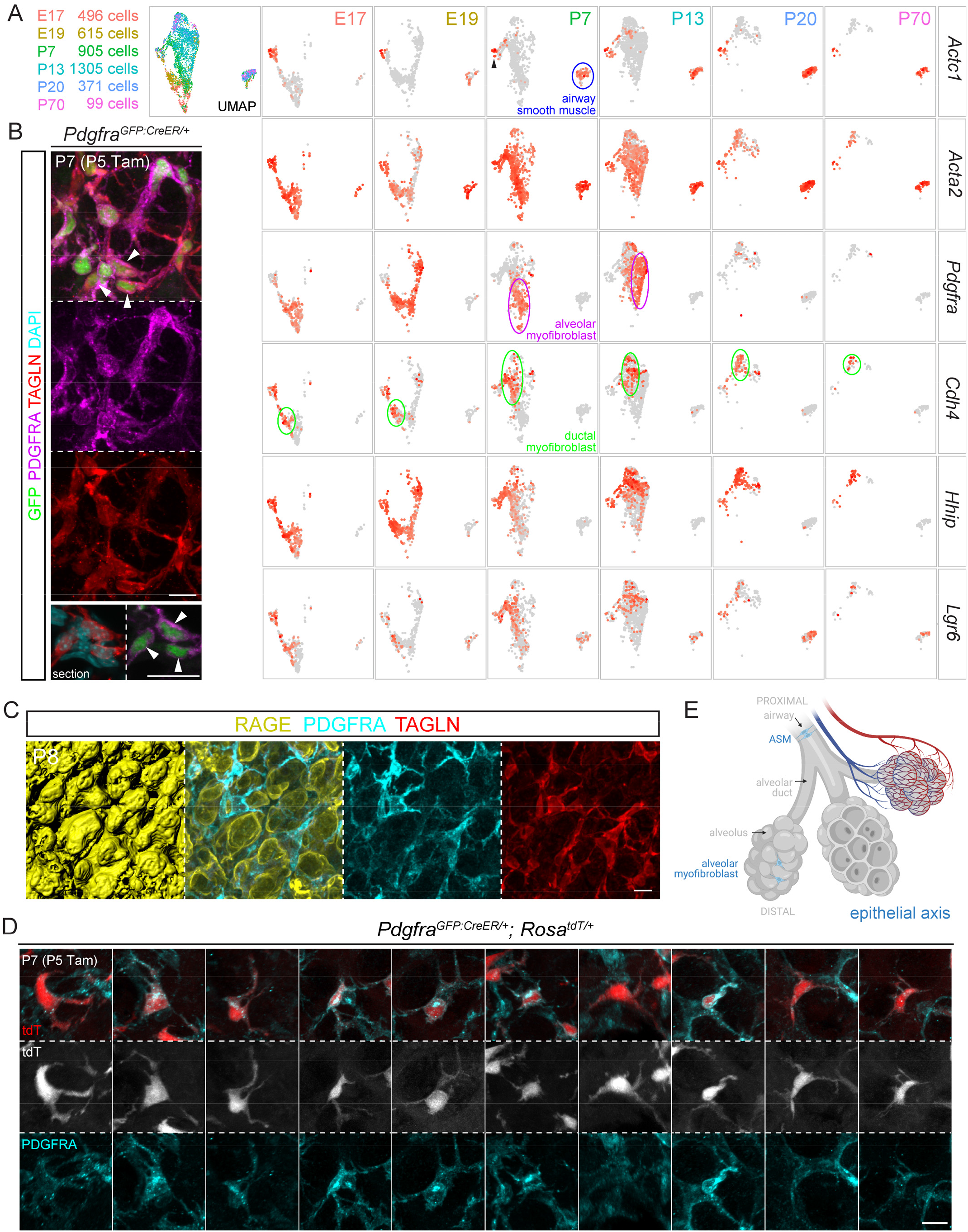
The epithelial axis includes airway smooth muscle cells, ductal myofibroblasts, and alveolar myofibroblasts, the last of which surround alveoli and disappear after classical alveologenesis. (**A**) ScRNA-seq UMAP and feature plots of lung mesenchymal cells of the epithelial axis, colored for developmental ages. Labeled for the P7 lung, airway smooth muscle cells form a distinct cluster, whereas the remaining cluster is also *Acta2*+ and contains a *Pdgfra*+ population (also present at P13 but absence at P20 and P70) and a *Cdh4/Hhip/Lgr6*+ population (present at all time points), corresponding to alveolar and ductal myofibroblasts. A small *Actc1*+ group (arrowhead) is adjacent the myofibroblast cluster, possibly reflecting a transition from the proximal airway smooth muscle cells to the distal myofibroblasts. (**B**) Immunostaining images showing PDGFRA-high cells (arrowhead), marked by bright GFP from *Pdgfra^GFP:CreER^* and perinuclear PDGFRA, express a contractile protein TAGLN. GFP-dim cells are not visible in this imaging setting. Tam, 300 ug tamoxifen. Scale: 10 um. (**C**) Immunostaining images and surface rendering to show PDGFRA-high, TAGLN+ alveolar myofibroblasts situated within grooves of alveolar type 1 cells (RAGE). Scale: 10 um. (**D**) Single cell morphology of 10 alveolar myofibroblasts from sparse genetic labeling. Tam, 300 ug tamoxifen. Scale: 10 um. (**E**) Diagram showing ASM cells and alveolar myofibroblasts of the epithelial axis. Created with BioRender.com.

### PDGFRA-high myofibroblasts surround alveoli and disappear after classical alveologenesis

Similar to PDGFRB and NOTCH3 (Fig. S3A), PDGFRA staining was diffuse throughout the cell and also concentrated in a perinuclear compartment in the neonatal lung (Fig. 3B); this perinuclear PDGFRA was absent in the mature lung (Fig. S3B), suggesting at least two types of PDGFRA-expressing cells. Consistent with this possibility, a histone-GFP knock-in allele of *Pdgfra* had identified GFP-bright and GFP-dim populations (Endale et al., 2017), which we hypothesized to correspond to those with and without perinuclear PDGFRA staining. As the level of histone-GFP would be confounded by the rate of cell division, we resorted to a GFP:CreER fusion knock-in allele of *Pdgfra* (Miwa and Era, 2015) and found that tamoxifen-induced nuclear GFP:CreER was high in cells with perinuclear PDGFRA, which accumulated TAGLN and thus were myofibroblasts (Fig. 3B). Unlike TAGLN, a commonly used marker ACTA2 aligned with PDGFRA cell extensions but did not reliably outline the nucleus for cell counting (Fig. S3C).

These perinuclear-PDGFRA contractile cells occupied grooves over the folding alveolar surface during classical alveologenesis (P3-P14) (Vila Ellis and Chen, 2020) and thus corresponded to alveolar myofibroblasts (Fig. 3C). Single-cell imaging using the *Rosa^tdT^* reporter showed frequently a strained tri-projection morphology and that each cellular projection was nucleus-length such that multiple cells were expected to assemble a network to constrain alveolar outpocketing (Fig. 3D, E).

### CDH4/HHIP/Lgr6 myofibroblasts surround alveolar ducts and persist in the adult lung

As predicted by scRNA-seq (Fig. 3A), CDH4+ myofibroblasts and PDGFRA-high myofibroblasts coexisted in neonatal lungs and constituted the TAGLN+ contractile cells (Fig. 4A). Wholemount immunostaining revealed in a Z-plane view a striking zonation of distal (i.e. surface) PDGFRA and proximal CDH4 (Fig. 4B). The proximal CDH4 zone corresponded to the alveolar ducts – tubular extensions of the airways but bordered by alveolar type 1 and type 2 cells. Arising from branch stalks as the airways did, alveolar ducts had wider airspace than the surrounding alveoli and could be best identified as they extended toward the lateral edge, instead of the lobe surface where tissue geometry made tubes less recognizable as they were shorter and interrupted by branching (Vila Ellis and Chen, 2020; Yang and Chen, 2014). We also found an HHIP antibody that marked cellular projections that largely aligned with CDH4, but also perinuclear regions for cell enumeration (Fig. 4C). As predicted from the Z-plane view (Fig. 4B), CDH4/HHIP cells specifically wrapped alveolar ducts, reminiscent of ASM cells wrapping the airways, albeit with larger gaps due to interruption by alveolar outgrowth (Fig. 4C). This continuation of CDH4/HHIP cells with ASM cells was supported and better visualized by the labeling of both cell populations by *Crh^Cre^* (Taniguchi et al., 2011), which we found in a screen for new mesenchymal cell drivers (Fig. S4). Notably, *Crh^Cre^* did not label other mesenchymal cells such as the VSM cells, supporting our model of three discrete mesenchymal axes. Similarly, an *Lgr6^GFP:CreER^* driver (Snippert et al., 2010), as predicted by scRNA-seq (Fig. 3A), labeled ACTA2+ contractile cells around airways and alveolar ducts – the latter of which were marked by HHIP – but not vessels (Fig. 4D). These cells that were labeled in the neonatal lung persisted in the mature lung (Fig. 4D).

**Figure 4:**
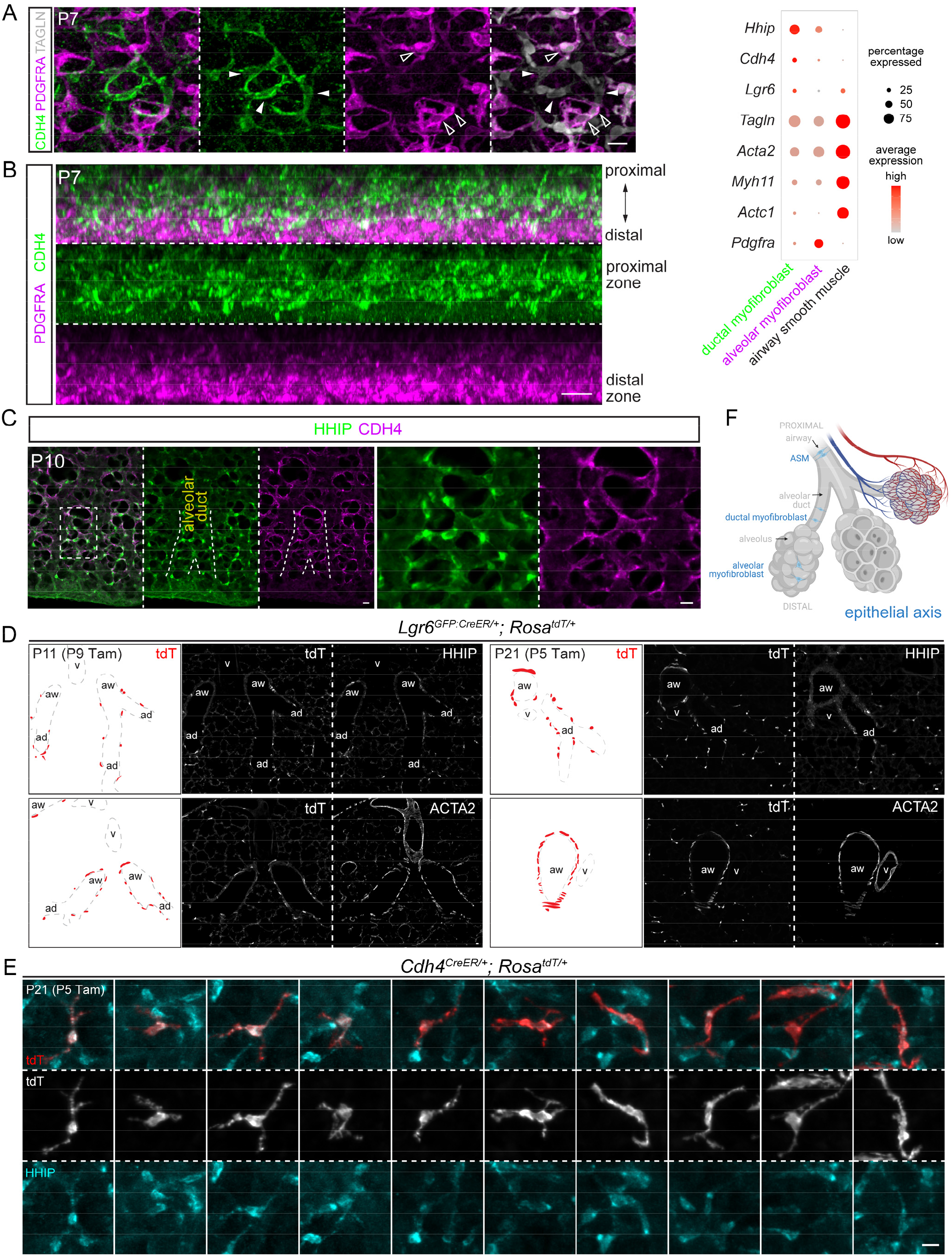
CDH4/HHIP/Lgr6 myofibroblasts surround alveolar ducts and persist in the adult lung. (**A**) Immunostaining images showing TAGLN+ contractile cells include PDGFRA+ and CDH4+ populations, as supported by the dot plot that includes additional markers. (**B**) Z-plane view of immunostaining images showing CDH4 and PDGFRA occupy the proximal and distal zones, corresponding to alveolar ducts and alveoli, respectively. Scale: 10 um. (**C**) Immunostaining images showing that HHIP and CDH4 respectively mark the perinuclear region and projections of cells around alveolar ducts. Scale: 10 um. (**D**) Immunostaining images and diagrams to show that *Lgr6^GFP:CreER^* labels airway (aw) smooth muscle cells (ACTA2+) and alveolar duct (ad) myofibroblasts (HHIP+), but not vascular (v) smooth muscle cells (ACTA2+). Labeled cells in neonatal lungs persist in the mature lung. Tam, 500 ug tamoxifen. Scale: 10 um. (**E**) Single cell morphology of 10 ductal myofibroblasts from sparse genetic labeling. Tam, 500 ug tamoxifen. Scale: 10 um. (**F**) Diagram showing ASM cells, ductal myofibroblasts, and alveolar myofibroblasts of the epithelial axis. Created with BioRender.com.

We additionally characterized a *Cdh4^CreER^* driver that was generated to label retinal neurons (Rousso et al., 2016) and found it, despite its inefficiency, specific to CDH4/HHIP+ cells (Fig. S5). The resulting sparse cell labeling showed strained cellular projections that were similar to those of PDGFRA-high myofibroblasts, but longer and often bidirectional, as expected for cells constraining a ductal structure wider than a spherical alveolus (Fig. 4E; note the different scale bar from that in Fig. 3D). In addition, *Cdh4^CreER^* lineage-labeled cells during the neonatal stage remained in the mature lung (Fig. 4E), consistent with persistence of these cells in our scRNA-seq data (Fig. 3A). Taken together, our data supported two populations of myofibroblasts – CDH4/HHIP/LGR6 ductal myofibroblasts and PDGFRA-high alveolar myofibroblasts – that are spatially separated, respectively persist or disappear after classical alveologenesis, and together with ASM cells form the proximal-distal epithelial axis, constraining the associated airway, alveolar duct, and alveolar epithelium (Fig. 4F). As observed in the gradual transition between VSM cells and pericytes, the epithelial axis also has intermediate cells as evidenced by *Actc1* expression in the myofibroblast clusters (Fig. 3A) as well as PDGFRA and CDH4/HHIP double positive cells (Fig. S3D).

### The interstitial axis, visualized by a new marker MEOX2, includes distal *Wnt2*+ PDGFRA-low cells and proximal IL33/DNER/PI16+ cells

Similarly, cells of the interstitial axis were reclustered into the aforementioned proximal *Twist2*+ (no reliable antibody identified) and distal *Wnt2*+ populations, the latter of which showed additional heterogeneity between developing and mature lungs – similar to that between immature and mature pericytes (Fig. 5A, 2A, Table S4). Supporting that the interstitial axis could be as discrete an entity as the vascular and epithelial axes, all interstitial mesenchymal cells specifically expressed *Meox2* – a nuclear protein that would also allow unequivocal cell identification and counting (Fig. S6A). We validated a MEOX2 antibody based on its expected absence in immune, epithelial, and endothelial cells, as well as PDGFRA-high alveolar myofibroblasts and PDGFRB pericytes (Fig. 5B, C). Instead, MEOX2 was in PDGFRA-low cells in the alveolar region, identifiable by lack of perinuclear PDGFRA staining and dim GFP from the *Pdgfra^GFP:CreER^* allele (Fig. 5D), consistent from prior studies using a histone-GFP reporter of *Pdgfra* (Endale et al., 2017). MEOX2+ cells were labeled by *Pdgfra^GFP:CreER^* with a high dose of the inducer tamoxifen, and had long wavy multidirectional protrusions occupying interstitial space (Fig. 5E, F). MEOX2 cells were in the vicinity of alveolar type 2 (AT2) cells, but not noticeably closer than other cell nuclei (Fig. S6B). Although a subset of AT2 cells might be particularly juxtaposed with MEOX2 cells, neither scRNA-seq nor scATAC-seq identified a distinct AT2 cell population in the mouse lung (Little et al., 2021). Oil red staining, commonly used to detect lipofibroblasts that were expected to corresponded to MEOX2+ cells as discussed earlier, was not specific to a particular cell type in neonatal or mature lungs (Fig. S6C).

**Figure 5:**
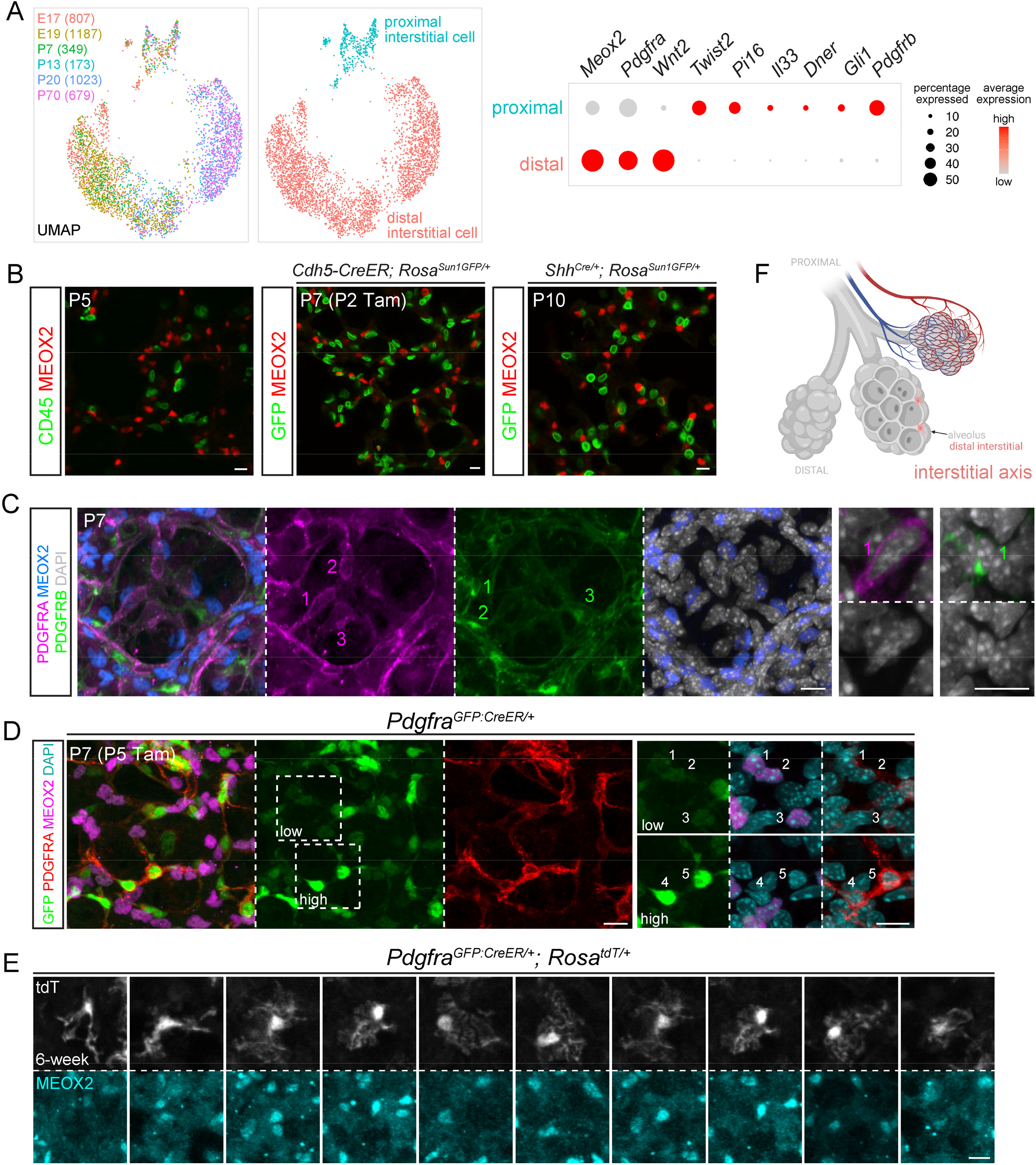
The interstitial axis is marked by MEOX2 and includes distal *Wnt2*-expressing, PDGFRA-low cells. (**A**) ScRNA-seq UMAPs of lung mesenchymal cells of the interstitial axis (cell number in parenthesis), colored by developmental ages (left) or cell types, as supported by relevant markers the dot plot. *Meox2* is lower in a subset of proximal interstitial cells. Distal interstitial cells form subclusters of immature (E17, E19, P7, P13) and mature (P20, P70) cells. (**B**) Immunostaining images showing MEOX2 is not expressed by immune cells (CD45) or genetically labeled endothelial *(Cdh5-CreER)* or epithelial *(Shh^Cre^)* cells. Genetic labeling is necessary to circumvent costaining of antibodies from the same species. Tam, 250 ug tamoxifen. Scale: 10 um. (**C**) Immunostaining images showing MEOX2 is not expressed by PDGFRA-high alveolar myofibroblasts or PDGFRB+ pericytes. Close-ups show perinuclear distribution of PDGFRA and PDGFRB. Scale: 10 um. (**D**) Immunostaining images showing that PDGFRA-low cells (#1-3) have dim GFP and express MEOX2, while PDGFRA-high cells (#4-5) have bright GFP and perinuclear PDGFRA. Tam, 300 ug tamoxifen. Scale: 10 um. (**E**) Single cell morphology of 10 distal interstitial cells from sparse genetic labeling with 3 mg tamoxifen 2 days before lung harvest. Scale: 10 um. (**F**) Diagram showing distal interstitial cells. Created with BioRender.com.

MEOX2 cells were also present more proximally, expressing low PDGFRA – identifiable by dim GFP from *Pdgfra^GFP:CreER^* – within bronchovascular bundles (Fig. 6A, B). To further characterize these proximal interstitial cells, we resorted to immunostaining for additional markers identified from scRNA-seq including *Il33*, *Dner*, and *Pi16*, as RNA probes were less reliable for low abundance genes and their speckly staining uninformative for discerning cell morphology. Consistent with prior studies (Dahlgren et al., 2019; Tsukui et al., 2020), IL33 was nuclear and expressed distally only by AT2 cells and proximally only by MEOX2 cells (Fig. 6C, S1C, S7A). Both DNER and PI16 marked the spindly perinuclear portion that tapered into thin processes (Fig. 6C, S7B). This cell morphology was best visualized using the *Pdgfrb^CreER^* driver, which scRNA-seq predicted to be active in both pericytes and proximal interstitial cells (Fig. 2A, 5A, S6A). Intriguingly, these elongated proximal interstitial cells were wedged between and basal to ASM cells (Fig. S7B, 6D). Notwithstanding occasional PDGFRA-low cells without MEOX2 due to possibly additional heterogeneity among the proximal interstitial cells (Fig. 6C), these data supported a distinct interstitial axis marked by MEOX2 and situating between vascular and epithelial trees (Fig. 6E).

**Figure 6:**
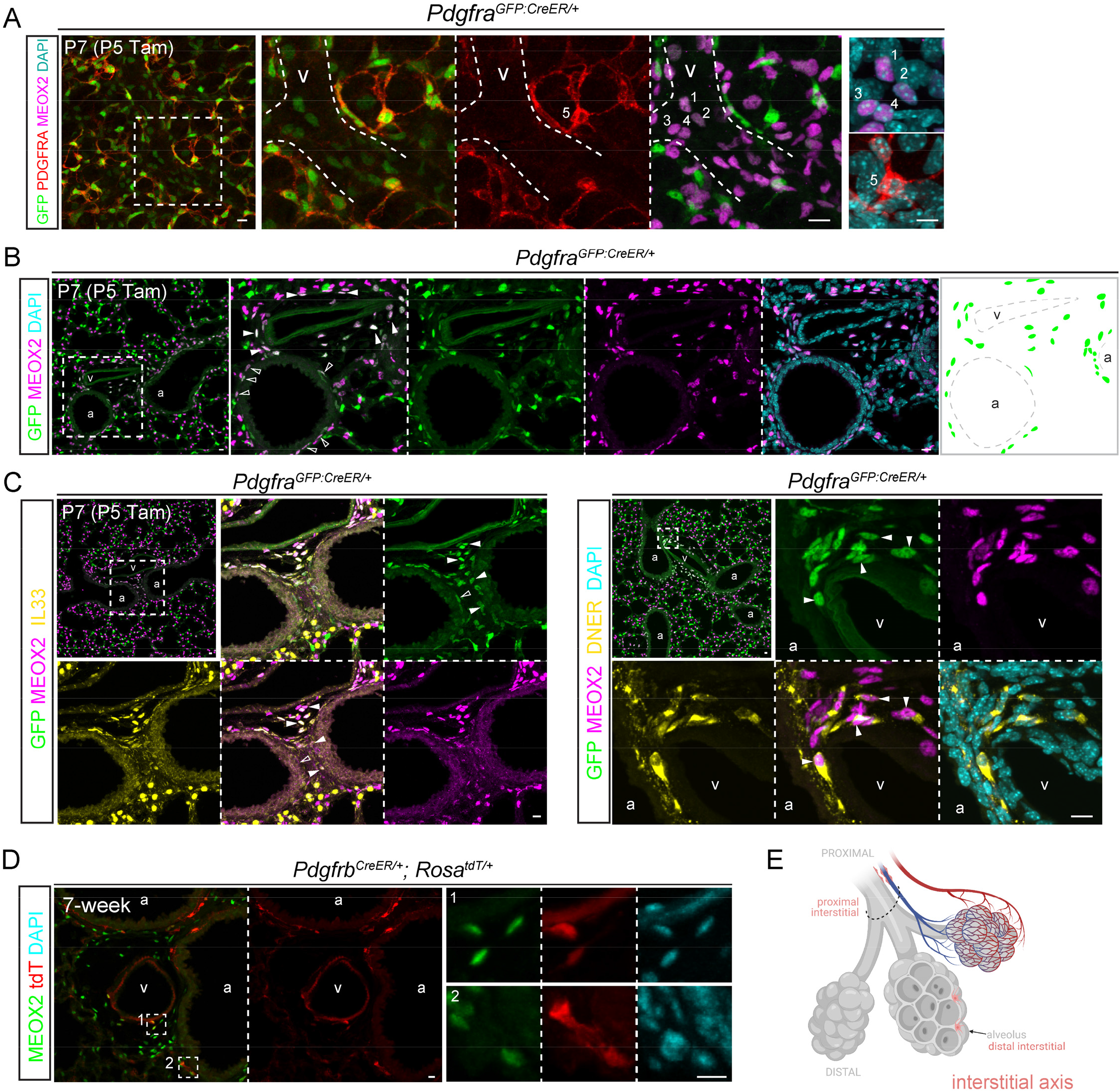
Proximal interstitial cells are in bronchovascular bundles and express MEOX2/IL33/DNER. (**A**) Immunostaining images showing dim GFP, MEOX2+ cells (#1-4) around a vessel (v) and bright GFP, PDGFRA-high cells (#5). Tam, 300 ug tamoxifen, which facilitates nuclear accumulation and hence detection of the GFP:CreER fusion protein. Scale: 10 um. (**B**) Immunostaining images and diagram showing GFP+, MEOX2+ cells around airways (a; open arrowhead) and vessels (v; filled arrowhead) within the bronchovascular bundle. Tam, 300 ug. Scale: 10 um. (**C**) Immunostaining images showing that GFP+, MEOX2+ cells (filled arrowhead) are IL33+ (left) and DNER+ (right) within bronchovascular bundles (a, airway; v, vessel). Occasional GFP+ cells are MEOX2- (open arrowhead). Tam, 300 ug. Scale: 10 um. (**D**) Immunostaining images showing that besides vascular smooth muscle cells, *Pdgfrb^CreER^* labels MEOX2+ cells within bronchovascular bundles (a, airway; v, vessel). Tam, 3 mg tamoxifen was administrated 2 days before lung harvest. Scale: 10 um. (**E**) Diagram showing proximal interstitial cells within the bronchovascular bundle (dashed semioval). Created with BioRender.com.

### Complementary lineage tracing supports apoptotic clearance of PDGFRA-high alveolar myofibroblasts

Within the distal compartment of the three axes, while developmental maturation was evident for pericytes and interstitial cells (Fig. 2A, 5A), the most striking change was the disappearance of PDGFRA-high alveolar myofibroblasts but persistence of CDH4/HHIP/LGR6 ductal myofibroblasts, as shown by *Lgr6^GFP:CreER^* and *Cdh4^CreER^* lineage tracing (Fig. 4D, E). Although myofibroblast clearance was recently reported (Hagan et al., 2020), we sought to clarify the distinct fates of the two myofibroblast populations, as well as genetic driver specificity using our newly identified molecular markers including MEOX2.

A contractile-gene driver *Myh11-CreER* (Wirth et al., 2008) marked alveolar myofibroblasts (99.5% efficiency from 1799 cells in 3 mice; Table S5), ductal myofibroblasts (98.2% efficiency from 425 cells in 3 mice), and unexpectedly pericytes (99.7% efficiency from 1530 cells in 3 mice) – perhaps reflecting their relatedness to VSM cells – but not MEOX2 interstitial cells in neonatal lungs (Fig. 7A, S8A). Tracing these labeled neonatal cells to the mature stage when alveolar myofibroblasts disappeared showed labeling in ductal myofibroblasts (98.4% efficiency from 290 cells in 3 mice) and pericytes (98.4% efficiency from 1501 cells in 3 mice), but still not MEOX2 cells, supporting that alveolar myofibroblasts does not become interstitial cells (Fig. 7A, S8A). To rule out the possibility that alveolar myofibroblasts became pericytes, we lineage-traced *Pdgfra^GFP:CreER^* cells and found that in the neonatal lung, *Pdgfra^GFP:CreER^* labeled mostly alveolar myofibroblasts (99.5% efficiency from 660 cells in 3 mice) and also MEOX2 interstitial cells (initially 3.4% efficiency from 873 cells in 3 mice, which increased over the tracing time to 59.6% efficiency from 1453 cells in 3 mice); none of them became pericytes in the mature lung (Fig. 7B, S8B). Consistent with pericytes being a distinct lineage, *Pdgfrb^CreER^* labeled cells remained pericytes in the distal compartment of both neonatal (98.5% efficiency from 416 cells in 3 mice) and mature (93.9% efficiency from 1037 cells in 3 mice) lungs (Fig. 7C). Notably, both *Myh11-CreER* and *Pdgfra^GFPCreER^* labeled cells, which overlapped for alveolar myofibroblasts, were found to express cleaved-Caspase 3 – consistent with apoptosis (Fig. 7D). These data were consistent with the notion that PDGFRA-high alveolar myofibroblasts undergo developmental apoptosis whereas CDH4/HHIP/LGR6 ductal myofibroblasts persist (Fig. 7E).

**Figure 7:**
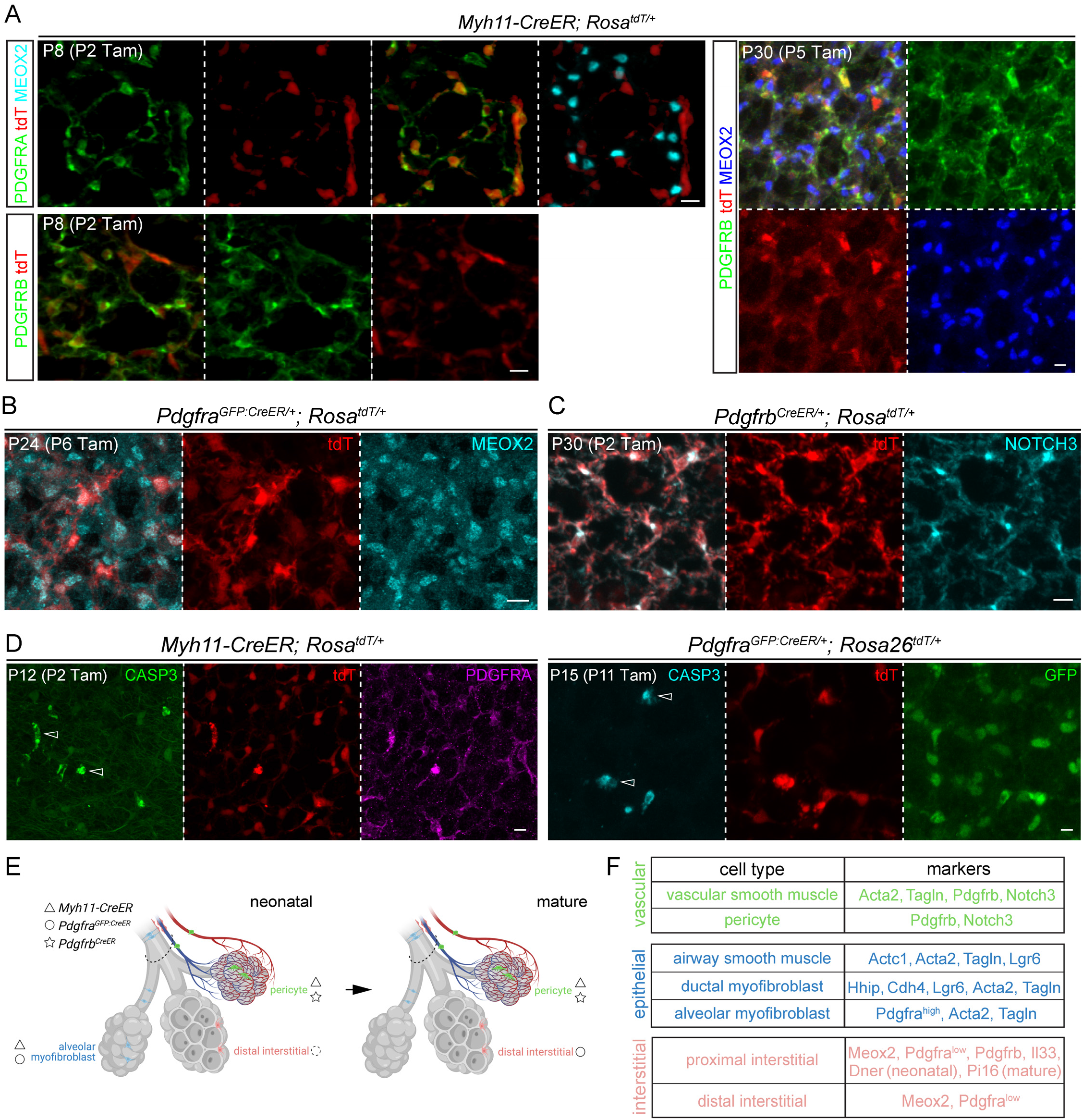
Complementary lineage tracing supports apoptotic clearance of PDGFRA-high alveolar myofibroblasts. (**A**) Immunostaining images showing that *Myh11-CreER* labels alveolar myofibroblasts (PDGFRA-high) and pericytes (PDGFRB+), but not distal interstitial cells (MEOX2+) in neonatal (P8) lungs. Pericytes remain labeled after tracing and distal interstitial cells remain unlabeled in mature (P30) lungs. Tam, 300 ug tamoxifen. Scale: 10 um. (**B**) Immunostaining images showing that *Pdgfra^GFP:CreER^* lineage-traced cells are MEOX2+. Tam, 300 ug tamoxifen. Scale: 10 um. (**C**) Immunostaining images showing that *Pdgfrb^CreER^* lineage-traced cells are pericytes (NOTCH3+). Tam, 250 ug tamoxifen. Scale: 10 um. (**D**) Immunostaining images showing that *Myh11-CreER* and *Pdgfra^GFP:CreER^* labeled cells are positive for cleaved Caspase 3 (CASP3; open arrowhead). Tam, 250 ug tamoxifen. Scale: 10 um. (**E**) Diagram showing labeled cells by the three drivers (symbols) in neonatal and mature lungs, consistent with developmental apoptosis of alveolar myofibroblasts. Dashed circle, inefficient labeling of distal interstitial cells, compared to alveolar myofibroblasts, by *Pdgfra^GFP:CreER^* in neonatal lungs. Created with BioRender.com. (**F**) Table summarizing the cell types of the three axes and their markers.

## DISCUSSION

In this study, we introduce a three-axis classification system for lung mesenchymal cells that integrates single-cell transcriptomic, spatiotemporal, morphological, and lineage information (Fig. 7F). Most intuitively, the vascular axis consists of proximal VSM cells and distal pericytes. The epithelial axis includes – from proximal to distal – ASM cells, hitherto unrecognized ductal myofibroblasts, and alveolar myofibroblasts, the last of which undergo developmental apoptosis. The interstitial axis shares a newly characterized marker MEOX2, fills the space between the vascular and epithelial trees such as that within the bronchovascular bundles, and includes proximal IL33+ cells and distal *Wnt2*+ cells – the latter of which has been called lipofibroblasts, albeit with no enrichment of the gene *Plin2.* This classification provides a framework to map the heterogeneous lung mesenchymal cells and predicts their functions and signaling interactions with nearby cell lineages, and can be extended to other organs.

Instead of the Cartesian system of XYZ coordinates, our three-axis classification has its root in the cylindrical coordinate system with an axial height for the proximal-distal location, a radial distance for the layered wrapping around a vascular or epithelial tube, and an azimuth for the cylindrical symmetry. This axial system is a natural way to characterize biology because tubes are fundamental building blocks and radial symmetry dates back to the primitive animal phylum of cnidarians. Locating mesenchyme cells with axial coordinates capitalizes on our better understanding of the vascular and epithelial trees, accounts for their likely supportive roles for each biological tree, and is consistent with their distinct developmental origins – as evidenced by the respective radial versus distal recruitment of VSM and ASM cells (Greif et al., 2012; Kumar et al., 2014), as well as the possible connection of the epithelial axis and progenitors or the interstitial axis to previously named subepithelial and submesothelial compartments, respectively (White et al., 2006). The resulting indirect demarcation of the proximal and distal compartments of the mesenchyme, possibly mediated by diffusible or mechanical signals, is expected to be more blurred than that for the endothelium or epithelium, leading to intermediate cells as observed for low-ACTA2 VSM cells, *Actc1*-expressing myofibroblasts, and PDGFRA/HHIP double positive cells (Fig. 2B, 3A, S3D). Such imprecision might also reflect intrinsic plasticity or mobility of mesenchymal cells to accommodate changes during tissue homeostasis and injury repair.

A notable prediction of the axial system that we have validated experimentally is the presence of ductal myofibroblasts associated with alveolar ducts, an epithelial structure connecting proximal airways with distal alveoli (Fig. 4). Like alveolar myofibroblasts, ductal myofibroblasts express contractile genes, have strained cellular projections, and cluster nearby on the UMAPs (Fig. 3, 4). Unlike alveolar myofibroblasts that undergo developmental apoptosis, ductal myofibroblasts persist in the mature lung, perhaps to corroborate or maintain the elastin cable scaffold (Wagner et al., 2015) or to serve as a main source of *Wnt5a* for nearby Wnt-responsive cells (Nabhan et al., 2018; Zacharias et al., 2018). Intriguingly, genetic polymorphisms in HHIP, a marker of ductal myofibroblasts (Fig. 3A, 4), have been linked to chronic obstructive pulmonary disease (COPD) (Zhou et al., 2012). The distinct developmental fates of ductal and alveolar myofibroblasts could explain the remaining *Fgf18-*lineage cells, which are expected to include both myofibroblasts (Fig. 1B) (Hagan et al., 2020), and also highlight the heterogeneity within secondary crest myofibroblasts that include cells around embryonic branch stalks and thus future alveolar ducts (Li et al., 2015; Zepp et al., 2021). Future studied are needed to understand the differential regulation, fate, and function of the two myofibroblast populations.

As whole genome sequencing has reversed the classical flow from protein biochemistry to gene discovery, single-cell genomics has cataloged transcriptionally and epigenetically distinct cell states in search of spatial, morphological, and functional features that traditionally define a cell type. Compared to in situ hybridization, immunostaining validates the protein products of transcriptional markers from single-cell genomics and reveals cell morphology in 3D tissues, but could be challenging to interpret due to distinct subcellular localization of protein markers, especially for intertwined lung cells. Notably, colocalization of image pixels is not always due to coexpression within the same cell, as exemplified by under-the-diffraction-limit juxtaposition and thus apparent colocalization of alveolar type 1 cell membrane markers such as AQP5 or RAGE with endothelial cell membrane markers such as ICAM2 or EMCN. On the other hand, coexpressed markers often do not colocalize, as exemplified by the nuclear localized NKX2-1 and junctional E-Cadherin that are expectedly present in every lung epithelial cell. Due to the complex morphology of lung mesenchymal cells (Fig. 2C, 3D, 4E, 5E), it is paramount to pinpoint the nucleus to localize and enumerate the cell populations identified in single-cell genomics. Accordingly, we identify a new marker MEOX2 for interstitial cells, complement ACTA2 with TALGN and CDH4 with HHIP, and characterize the perinuclear accumulation of PDGFRA and PDGFRB – with the added benefit of using perinuclear PDGFRA to distinguish PDGFRA-high versus low cells.

This battery of molecular tools shed light on lineage-tracing experiments. For example, *Pdgfra^GFP:CreER^* labels alveolar myofibroblasts and, less efficiently due to the lower *Pdgfra* expression, interstitial MEOX2 cells (Fig. 5, 6, 7), whereas *Pdgfrb^CreER^* labels pericytes and proximal interstitial cells (Fig. 2, 6) that are also referred to as adventitial fibroblasts (Tsukui et al., 2020), which might give rise to pathological myofibroblasts (Rock et al., 2011; Tsukui et al., 2020) and might also be the *Gli1^CreER^*-lineage cells promoting airway epithelium repair (Cassandras et al., 2020; Moiseenko et al., 2020). While the reported Axin2+ Myofibrogenic Progenitors (AMPs) seemingly correspond to pericytes and the Mesenchymal Alveolar Niche Cells (MANCs) proximal MEOX2 cells based on their distributions on the UMAPs, future work is needed to clarify the additional heterogeneity as a result of Wnt-signaling, as reflected by the apparent stochastic expression of *Axin2* (Fig. S1C), as well as enriched expression of Wnt coreceptors *Lgr5* and *Lgr6* in cells of the epithelial axis (Fig. 3, 4). These molecular analyses, integrated with cell-type-specific morphology – analogous to Golgi staining of neuronal cell morphology (Vints et al., 2019) – and functional studies, will lead to a multifaceted definition of lung mesenchymal cell types.

## METHODS

### Mus musculus

The mouse protocols used for this research complied with the regulations of MD Anderson Cancer Center’s Institutional Animal Care and Use Committee. Wild type C57BL/J mice were used for validation of gene-based molecular markers with immunostaining. The mouse strains for lineage labeling and tracing experiments were *Crh^Cre^* (Taniguchi et al., 2011), *Pdgfra^GFP:CreER^* (Miwa and Era, 2015), *Cdh4^CreER^* (Rousso et al., 2016), *Myh11-CreER* ((Wirth et al., 2008), *Pdgfrb^CreER^* (Cuervo et al., 2017), *Lgr6^GFP:CreER^* (Snippert et al., 2010), *Shh^Cre^* (Harfe et al., 2004), *Cdh5-CreER* (Sorensen et al., 2009), *Rosa^Sun1GFP^* (Mo et al., 2015), and *Rosa^tdT^* (Madisen et al., 2010). Intraperitoneal injections of tamoxifen (T5648; Sigma) dissolved in corn oil (C8267; Sigma) were administered to induce Cre recombination; specific dosages are detailed in figure legends. P0-P12 was considered the neonatal stage.

### Antibodies

The following antibodies were used for immunofluorescence: goat anti-platelet derived growth factor-alpha (PDGFRA, 1:1000, AF1062, R&D Systems), rat anti-PDGFRA (PDGFRA, 1:1000, 14-1401-82, eBioscience), rat anti-cadherin-4 (CDH4, 1:20, MRCD5, Developmental Studies Hybridoma Bank), rat anti-platelet derived growth factor receptor-beta (PDGFRB, 1:1000, 14-1402-82, eBioscience), goat anti-platelet derived growth factor receptor-beta (PDGFRB, 1:1000, AF1042, R&D Systems), goat anti-notch3 (NOTCH3, 1:500, AF1308, R&D Systems), rabbit anti-mesenchyme homeobox 2 (MEOX2, 1:1000, NBP2-30647, Novus Biological), rabbit anti-cleaved caspase-3 (CASP3, 1:500, 9661, Cell Signaling), goat anti-peptidase inhibitor 16 (PI16, 1:1000, AF4929, R&D Systems), rat anti-protein tyrosine phosphatase, receptor type, C (CD45, 1:2000, 14-0451-81, eBioscience), Alexa488-conjugated mouse anti-smooth muscle actin (ACTA2, 1:1000, Santa Cruz Biotechnology, sc-32251 AF488), Cy3-conjugated mouse anti-smooth muscle actin (ACTA2, 1:1000, Sigma, C6198), Alexa647-conjugated mouse anti-smooth muscle actin (ACTA2, 1:1000, Santa Cruz Biotechnology, sc-32251 AF647), rat anti-advanced glycosylation end product-specific receptor (RAGE, 1:1000, R&D Systems, MAB1179), rabbit anti-transgelin (TAGLN, also known as SM22, 1:1000, Abcam, ab14106), chicken anti-green fluorescent protein (GFP, 1:5000, Abcam, AB13970), goat anti-hedgehog-interacting protein (HHIP, 1:1000, R&D Systems, AF1568), guinea pig anti-lysosomal-associated membrane protein 3 (LAMP3, 1:500, Synaptic Systems, 391005), mouse anti-NK2 homeobox 1 (NKX2-1, 1:500, Leica Biosystems, TTF-1-L-CE), goat anti-interleukin 33 (IL33, 1:500, R&D Systems, AF3626), rabbit anti-calponin (CNN1, 1:1000, Abcam, ab46794), and goat anti-delta/notch-like EGF repeat containing (DNER, 1:1000, AF2264, R&D Systems).

### Cell dissociation and FACS

The experiment protocol for cell dissociation and fluorescence-activating cell sorting (FACS) was performed as described (Vila Ellis et al., 2020). After perfusion through the right heart ventricle with PBS, the whole lung was removed, separated into lobes in PBS and collected in ice-cold Eppendorf tubes with RPMI (ThermoFisher, 11875093) or Lebovitz’s Media (Gibco, 21083027). Lung lobes were minced with forceps and digested with 2 mg/mL Collagenase Type I (Worthington, CLS-1, LS004197), 2 mg/mL Elastase (Worthington, ESL, LS002294), and 0.5 mg/mL DNase I (Worthington, D, LS002007) for 30 min at 37°C using a dry block incubator. The tissue was resuspended at 15 minutes of incubation. The enzymatic digestion was stopped at 30 minutes by adding fetal bovine serum (FBS, Invitrogen, 10082-139) to a 20% concentration.

Samples were placed on ice and the following steps were performed in the cold room. The samples were resuspended until homogeneous, filtered with a 70 μm Falcon cell strainer (Falcon, 352350) and centrifuged in 2 mL tubes for 1 minute at 5000 rounds per minute (RPM). After supernatant removal, 1 mL red blood cell lysis buffer (15 mM NH4Cl, 12 mM NaHCO3, 0.1 mM EDTA, pH 8.0) was added gently to the sample, followed by a wait time of 3 minutes on ice and centrifugation. After performing this step twice, cells were washed and resuspended with 1 mL of icecold Lebovitz’s media supplemented with10% FBS. Cells were filtered into a 5 mL glass tube with cell strainer cap (Falcon, 352235) and transferred to a 2 mL Eppendorf tube for immunostaining for 30 minutes on ice for the immune cells (CD45-PE/Cy7, 1:250, 103114, BioLegend), endothelial cells (ICAM2-A647, 1:250, A15452, Life Technologies), and epithelial cells (ECAD-488, 1:250, 53-3249-80, eBioscience). Cells were centrifuged, washed with Lebovitz/10%FBS, and filtered into a 5 mL glass tube with cell strainer cap. After incubating cells with SYTOX Blue (Invitrogen, S34857), cells were analyzed using BD FACSAria Fusion Cell Sorter to evaluate cell viability and gating strategy for epithelial, endothelial, immune and mesenchymal cells. CD45 negative cells were selected, from those ICAM2 negative cells were selected, and from those ECAD negative cells were selected, resulting in the triple negative mesenchymal cell population. An equal proportion of immune, endothelial, epithelial and mesenchymal lineage cells were collected into a single tube and concentrated through centrifugation at 4°C for 5 minutes at 5000 RPM. Supernatant that contained ambient RNA was removed, resuspended in 250 μL Lebovitz/10%FBS and used for 10X Genomics library preparation.

### Single-cell RNA-Seq

Sequencing and analysis were performed as described (Vila Ellis et al., 2020). Sorted samples were prepared for sequencing using the Chromium Single Cell Gene Expression Solution Platform (10X Genomics) along with the Chromium Single Cell 3’ Library and Gel Bead Kit (V2, revD). Samples were sequenced with an Illumina NextSeq500 using a 26X124 sequencing run format with 8 bp index (Read1). Sequencing output files underwent pipeline analysis with “cellranger count” and “cellranger aggregate” that result in files for easy analysis with Loupe Cell Browser (10X Genomics) and Seurat 3.1, an R-package for quality control, normalization, and data exploration (https://satijalab.org/seurat/). Quality control metrics were analyzed per sample. Cells were filtered using the following criteria: no less than 200 genes, no more than 6000 transcripts, mitochondrial percentage up to 15% to preserve embryonic cells that are enriched in mitochondria. Additional normalization using the R package, harmony, reduced batch effects across samples (https://portals.broadinstitute.org/harmony/index.html). Clusters were analyzed with FindAllMarkers function to identify differentially expressed genes and markers for known and novel lung mesenchymal populations. Trajectory analysis was performed with Monocle 2.8 using the top 1500 genes (Qiu et al., 2017). R script is available as Supplemental File. Raw data were deposited at GEO under accession number GSE180822.

### Lung harvest

Lungs were processed as described in our previous publications (Little et al., 2019; Vila Ellis et al., 2020; Yang et al., 2016). Briefly, mice were anesthetized with Avertin (2, 2, 2-tribromoethanol) [T48402, Sigma]. After opening the abdominal cavity, the aorta was cut for exsanguination, followed with opening of the rib cage and perfusion of the heart’s right ventricle with PBS. The trachea was separated from posterior structures using blunt dissection and incised for canula insertion. Thread and a surgical knot were used to stabilize the canula to allow gravity drip lung inflation at 25 cm H_2_O pressure with 0.5% paraformaldehyde (PFA; P6148; Sigma) in PBS for 5 minutes. Lungs were collected in 0.5% PFA/PBS and incubated for 3 hours at room temperature on a rocker. After fixation, samples were washed again in PBS overnight at 4C for further processing.

### Section immunostaining

The immunostaining protocol was performed as described in our previous publications (Little et al., 2019; Vila Ellis et al., 2020; Yang et al., 2016). Fixed lungs were separated into lobes. The middle and caudal lobes were cryoprotected in 20% sucrose in PBS containing 10% optimal cutting temperature (OCT) embedding compound (4583; Tissue-Tek) overnight at 4°C for rapid dry-ice freezing and -80°C storage. Cryosections of 20 μm were cut and dried for 1 hour at room temperature. After hydrophobic rectangles were drawn on the sections with a PAP pen, the samples were placed in a humid chamber and washed with PBS 3 times for 5 minutes each. Tissues were blocked with diluted 5% donkey serum (017-000-121, Jackson ImmunoResearch) in PBS with 0.3% Triton X-100 (PBST) for 1 hour at room temperature, followed with primary antibody incubation overnight at 4°C. Next, lung tissue sections were washed in a coplin jar with PBS for 30 minutes at room temperature and PBS excess was removed with a vacuum. Tissues were incubated with secondary antibody mixture for 1-2 hr, followed with a 30 minute PBS wash in a coplin jar at room temperature. Sections were mounted with Aquamount (18606, Polysciences) and covered with 24 × 50 mm coverslip (No. 1, VWR). Samples were imaged using Nikon A1 Plus and Olympus FV1000 confocal microscopes.

For oil red staining, a stock solution of 0.25 g Oil Red O Sigma (O0625-25G) was dissolved in 50 mL of isopropanol. Lung tissue sections of 20 um thick were immunostained as above without normal donkey serum and post-fixed with 2% PFA for 1 hour. A working oil red solution was freshly prepared by diluting the stock with distilled deionized water (ddH2O) in a 6:4 ratio and let stand for 10 minutes before being filtered first with a 70 um Falcon cell strainer and then a 0.22 um pore-size filter membrane to remove oil red precipitate. Immunostained lung sections were washed with PBS for 5 minutes three times and incubated in working oil red solution for 30 minutes, followed by three 30-second washes in ddH2O and then washed under running tap water for 10 minutes.

### Whole mount immunostaining

The outer edge of cranial and left lung lobes was cut into strips and blocked with donkey serum for 1 hour. They were incubated with primary antibody mixture at 4°C in a rocker overnight, followed with 1 hour washes with PBS with 1% Triton X-100 and 1% Tween-20 (PBSTT) 3 times. Lung strips were incubated with secondary antibody mixture overnight at 4°C, followed with PBSTT washes as previously done for primary antibody incubation. Lung strips were then fixed using 2% PFA for 2-4 hours and washed with PBS 3 times for 5 minutes each. Samples were placed over on plain microscope slides (8201, Premier) with Aquamount and covered with 22 × 22 mm coverslips (No 1,48366 067, VWR). Immunostained whole lung lobes were imaged on an optical projection tomography scanner (Bioptonics, 3001M).

### Confocal imaging and processing

Samples were imaged with both microscopes using 4-color imaging (DAPI, GFP, TXR, Cy5) and 1-μm-step-size 20- to 50-μm stacks. Images were analyzed using Imaris 7.7.2 ^®^ (Bitplane, http://www.bitplane.com/imaris), ImageJ (U.S. National Institutes of Health, https://imagej.nih.gov/ij/) software. Quantification and was performed with Imaris^®^ software and GraphPad Prism^®^ software. Biological replicates and mouse number are specified within the figures.

## ACKNOWLEDGEMENT

We thank Dr. Margo P Cain and Belinda J Hernandez in our lab for generating scRNA-seq data for embryonic and P70 lungs. We thank Dalia Hassan, Celine SL Kong, and Andrés Gutiérrez for assistance with quantification of genetic drivers. The University of Texas MD Anderson Cancer Center DNA Analysis Facility and Flow Cytometry and Cellular Imaging Core Facility are supported by the Cancer Center Support Grant (CA #16672). This work was supported by the University of Texas MD Anderson Cancer Center Start-up and Retention Fund, and National Institutes of Health R01HL130129 and R01HL153511 (JC) and F31HL149232 (ONP).

## AUTHOR CONTRIBUTIONS

ONP and JC designed and performed research, and wrote the paper.

