## Supplemental Figures for "Three-axis classification of mouse lung mesenchymal cells reveals two populations of myofibroblasts"

**8 Supplemental Figures**

**5 Supplemental Tables**

**1 Supplemental File**

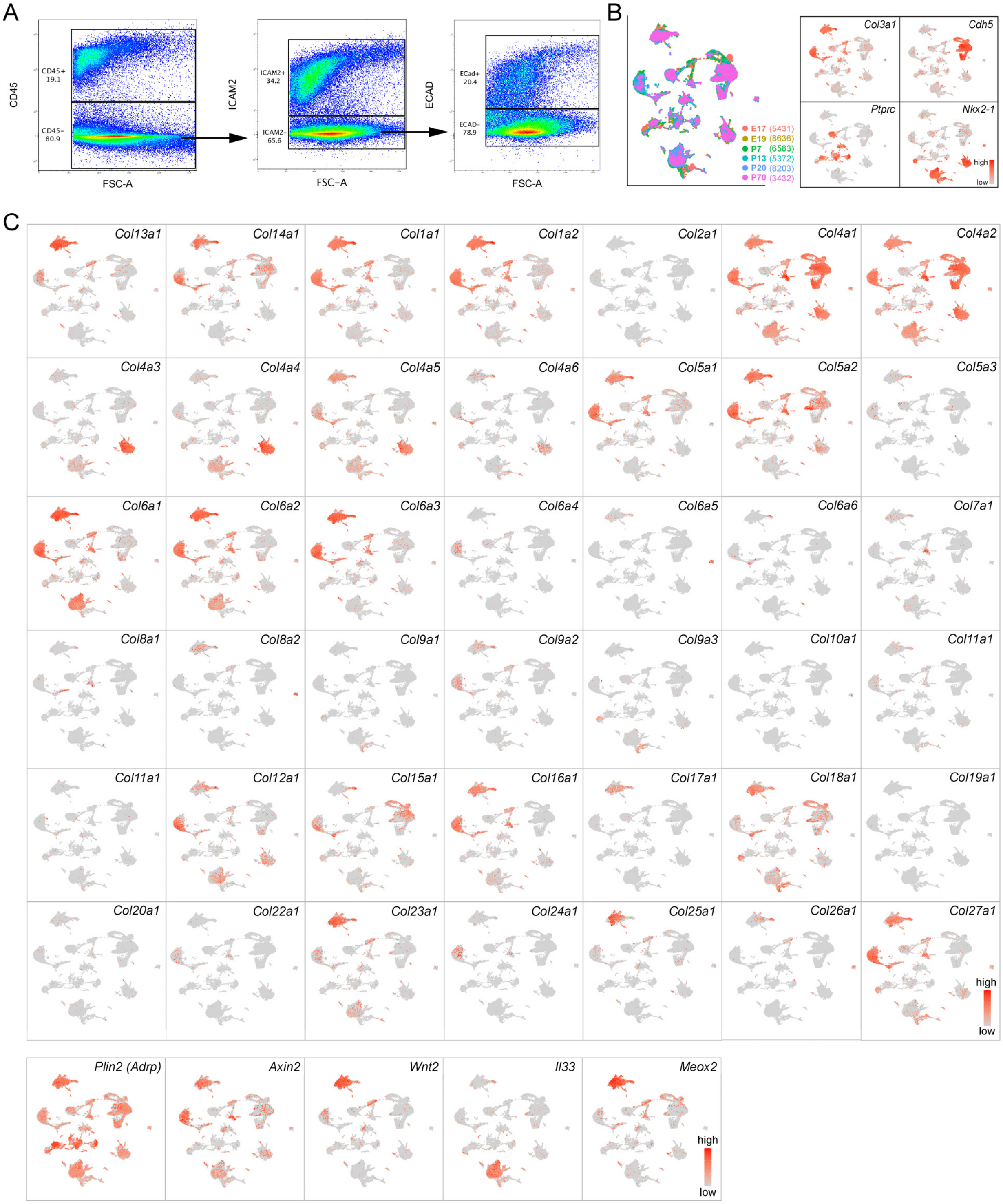


**Fig. S1: Single-cell RNA-seq of mouse lungs.**

(**A**) Representative FACS gating strategy to separate immune (CD45), endothelial (ICAM2), epithelial (ECAD), and mesenchymal (triple negative) cells.

(**B**) ScRNA-seq UMAP (cell number in parenthesis) and feature plots to identify mesenchymal (*Col3a1*), endothelial (*Cdh5*), immune (*Ptprc*), and epithelial (*Nkx2-1*) cells.

(**C**) ScRNA-seq feature plots showing matrix genes (top) in non-mesenchymal cells, wide-spread expression of a presumable lipofibroblast marker *Plin2* (also known as *Adrp*) and a Wnt-signaling target gene *Axin2*, and specific expression of *Wnt2*, *Il33*, and *Meox2* (bottom row).

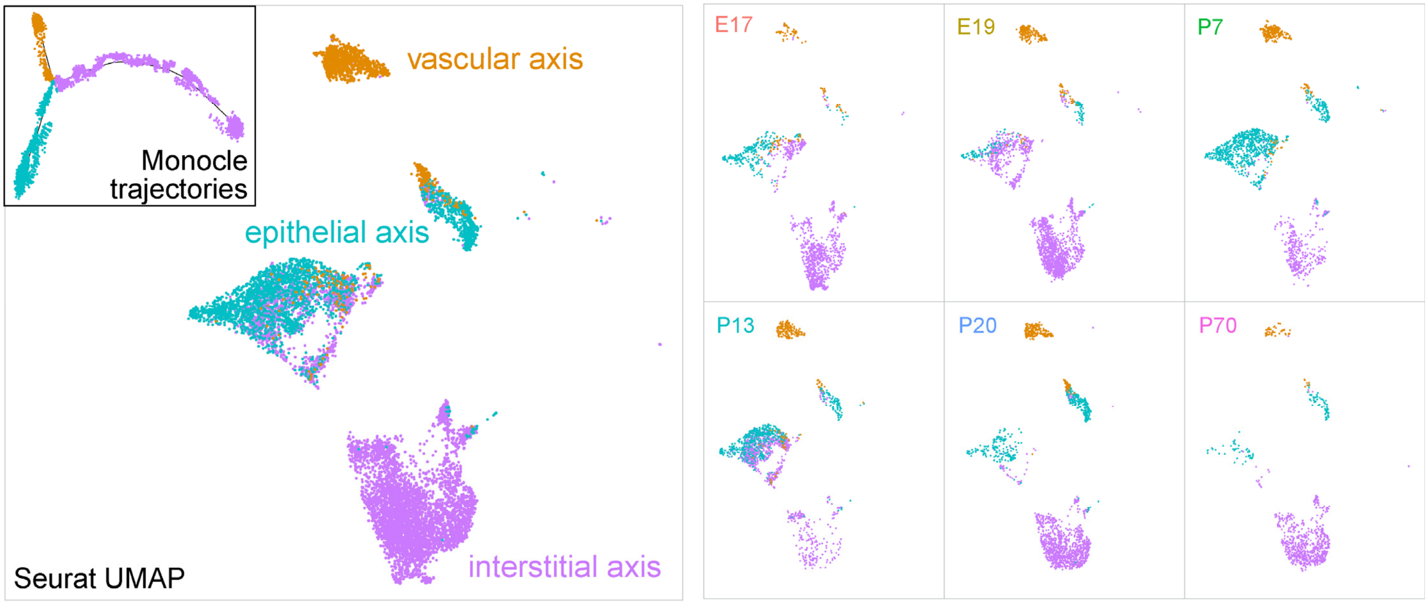


**Fig. S2: Distribution of Monocle trajectories on Seurat UMAPs.**

Cells in the 3 Monocle trajectories are colored on Seurat UMAPs corresponding to the vascular, epithelial, and interstitial axes.

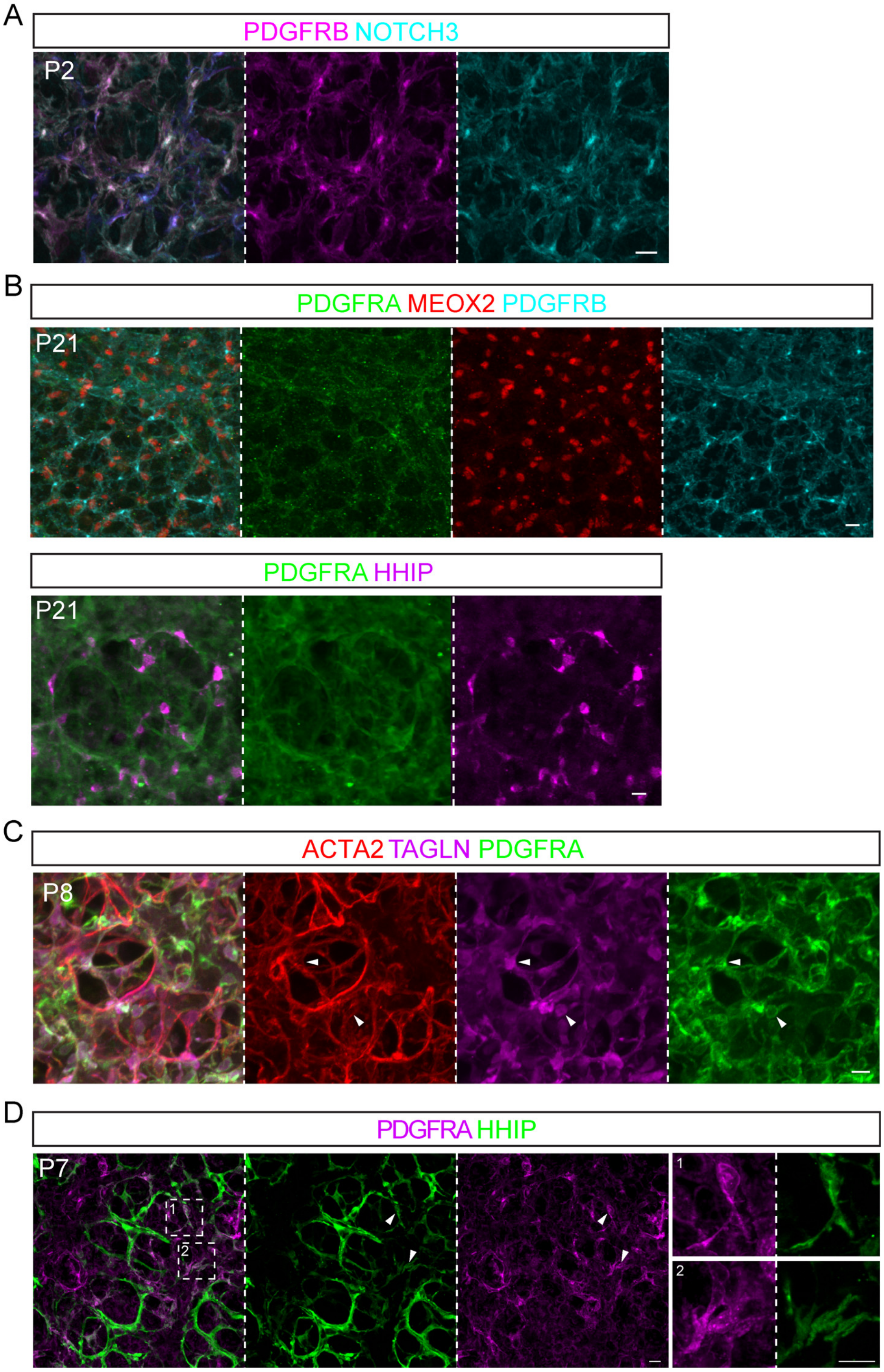


**Fig. S3: Further validation of mesenchymal cell markers.**

(**A**) PDGFRB and NOTCH3 are co-expressed in pericytes and have perinuclear accumulation. Scale: 10 um.

(**B**) PDGFRA staining in MEOX2+ cells in the mature lung is diffuse and has no perinuclear accumulation, as compared to perinuclear PDGFRB staining in pericytes (top) and perinuclear HHIP staining in ductal myofibroblasts (bottom). Scale: 10 um.

(**C**) ACTA2 staining does not reliably mark the cell nucleus, in contrast to TAGLN and PDGFRA (arrowhead). Scale: 10 um.

(**D**) Occasional PDGFRA and CDH4/HHIP double positive cells (arrowhead) are possibly intermediates between ductal and alveolar myofibroblasts. Scale: 10 um.

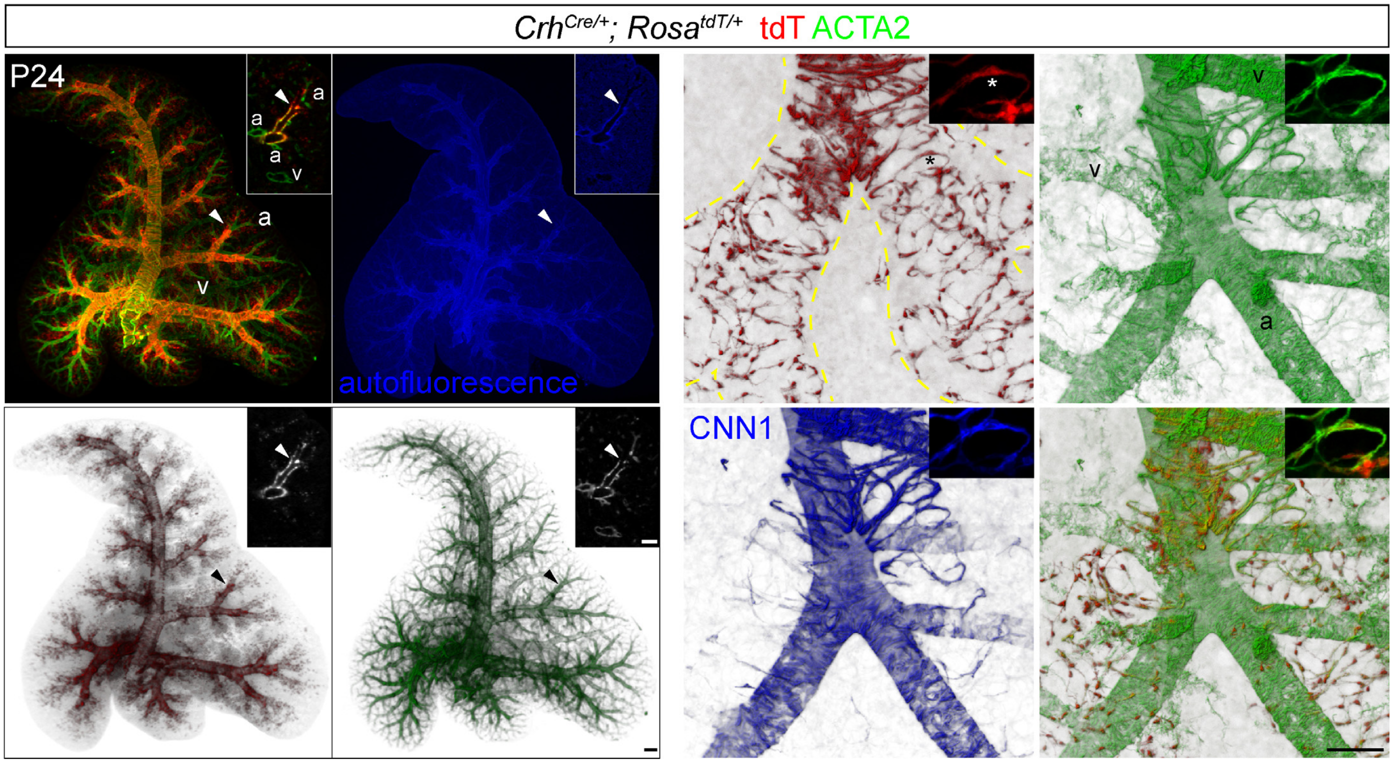


**Fig. S4: Characterization of a *Crh*$^{Cre}$ driver.**

Left: Optical projection tomography images showing that *Crh*$^{Cre}$ labeled cells surround airways and alveolar ducts (arrowhead, bronchoalveolar duct junction recognizable by the abrupt thinning of the epithelium autofluorescence), but not arterioles (a) and venules (v). Scale: 250 um.

Right: *Crh*$^{Cre}$ labeled ductal myofibroblasts extend beyond the airways and express a low level of contractile proteins ACTA2 and CNN1. Asterisk: nucleus of the cell-of-interest. Scale: 100 um.

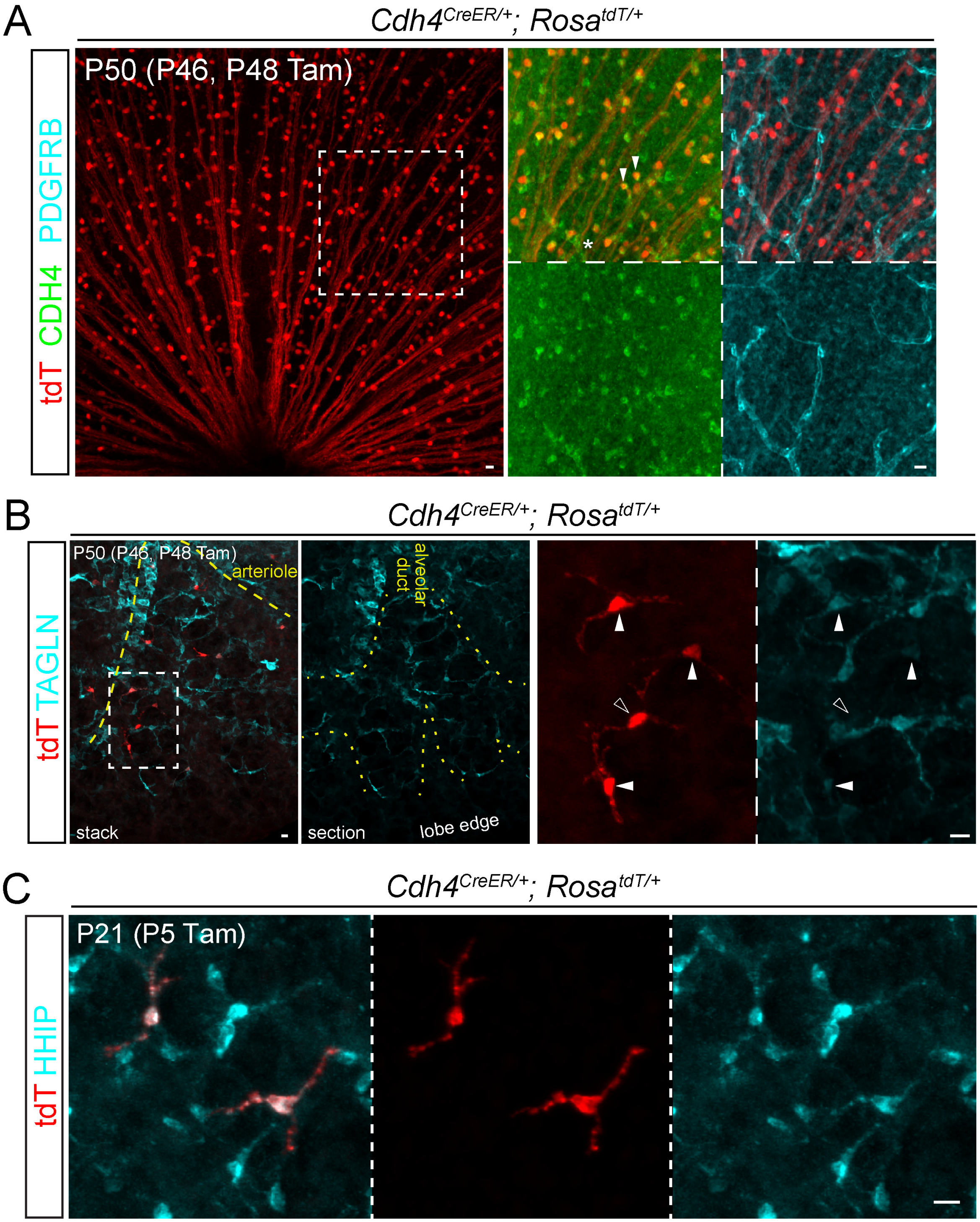


**Fig. S5: Characterization of a *Cdh4$^{CreER}$* driver.**

(**A**) Flat mount immunostained retina showing expected, efficient labeling of CDH4+ ganglion cells (filled arrowhead). Asterisk, CDH4 staining in some vessels marked by PDGFRB+ pericytes. Tam, 3 mg tamoxifen. Scale: 10 um.

(**B**) Inefficient but specific labeling of ductal myofibroblasts in the mature lung. Some ductal myofibroblasts have reduced TAGLN (filled versus open arrowhead). Tam, 3 mg tamoxifen. Scale: 10 um.

(**C**) Lineage-traced ductal myofibroblasts expressing HHIP. Tam, 500 ug tamoxifen. Scale: 10 um.

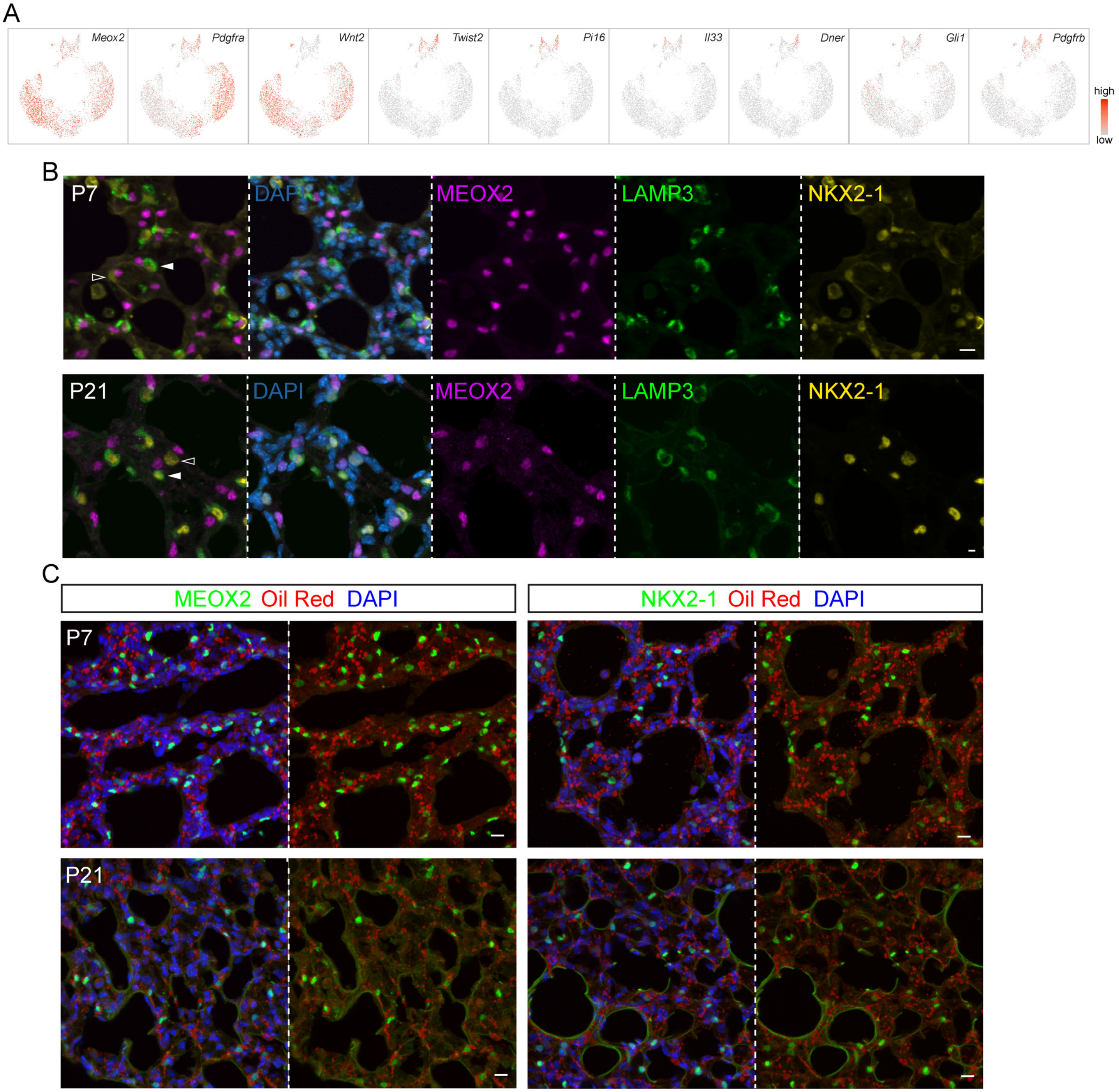


**Fig. S6: Localization of MEOX2+ interstitial cells and oil red stained cells.**

(**A**) ScRNA-seq feature plots of interstitial cells. See UMAP in Fig. 5A.

(**B**) Immunostaining images showing that MEOX2+ cells are not closer to alveolar type 2 cell nuclei (filled arrowhead; LAMP3+ NKX2-1+) than to alveolar type 1 cell nuclei (open arrowhead; LAMP3- NKX2-1+) or any other nuclei (DAPI). Scale: 10 um.

(**C**) Oil red stained lipid droplets are wide-spread and not specific to MEOX2+ interstitial cells or NKX2-1+ epithelial cells. Scale: 10 um.

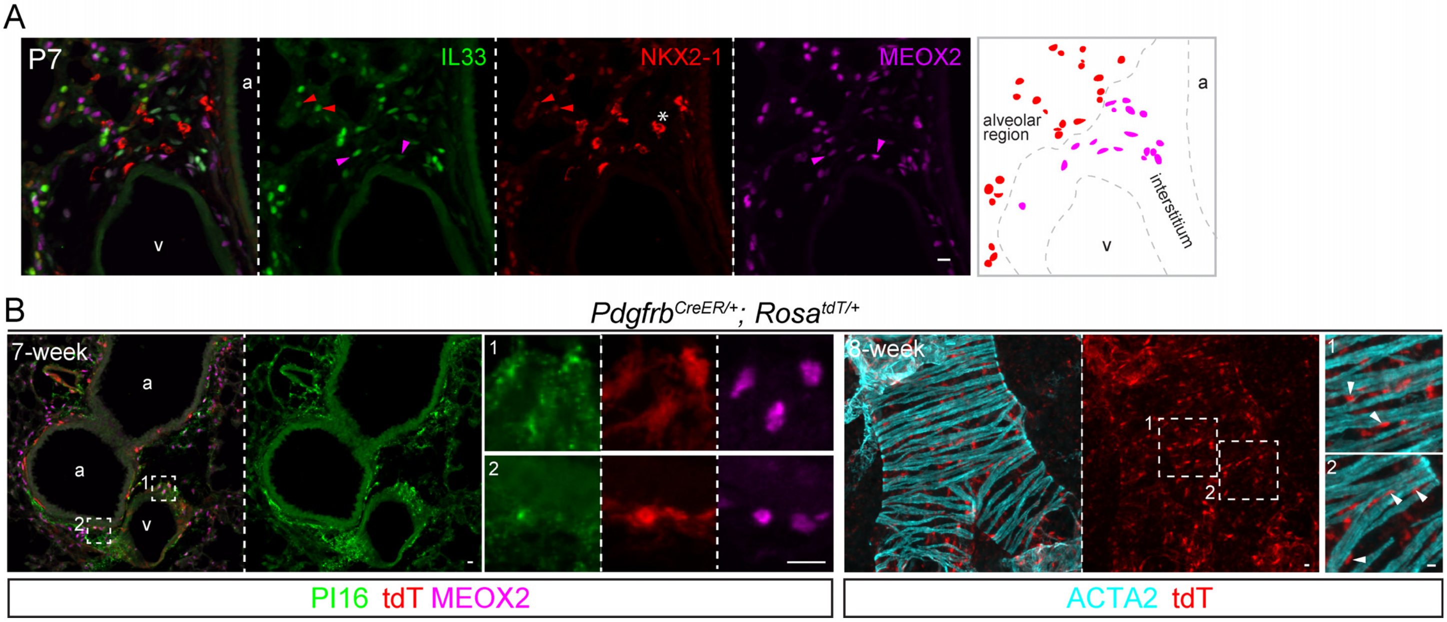


**Fig. S7: Further characterization of proximal MEOX2+ interstitial cells.**

(**A**) Immunostaining images and diagram showing that IL33 marks MEOX2+ cells in the bronchovascular bundle (a, airway; v, vessel), but NKX2-1 epithelial cells in the alveolar region. Asterisk, stained immune cells from the mouse NKX2-1 antibody. Scale: 10 um.

(**B**) Section (left) and wholemount (right) immunostaining images to show that *Pdgfrb*$^{CreER}$ labeled proximal interstitial cells express PI16 and MEOX2 and are between ACTA2+ airway smooth muscle cells (a, airway; v, vessel). 3 mg tamoxifen was administrated 48 hr before lung harvest. Scale: 10 um.

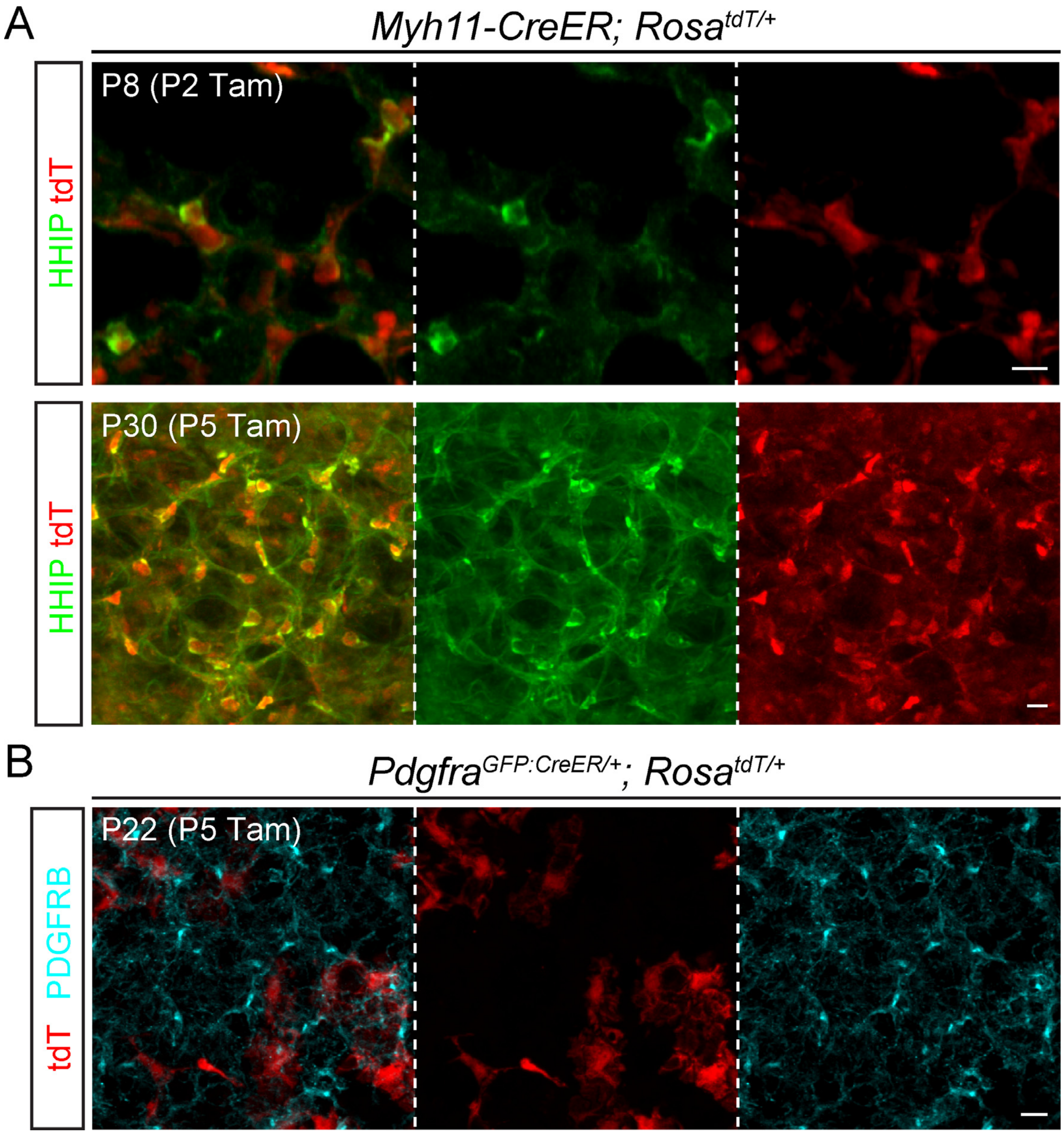


**Fig. S8: *Myh11-CreER* labels ductal myofibroblasts.**

(**A**) Immunostaining images showing *Myh11-CreER* labels HHIP+ ductal myofibroblasts in the neonatal lung (top), which persist in the mature lung (bottom). Tam, 300 ug tamoxifen. Scale: 10 um.

(**B**) Immunostaining images showing *Pdgfra*$^{GFP:CreER}$ labeled cells do not trace into pericytes (PDGFRB). Tam, 300 ug tamoxifen. Scale: 10 um.

**Table S1: Markers for the 24 clusters in Figure 1C.**

**Table S2: Markers for cell populations in the vascular axis in Figure 2A.**

**Table S3: Markers for cell populations in the epithelial axis in Figure 3A.**

**Table S4: Markers for cell populations in the interstitial axis in Figure 5A.**

**Table S5: Cell quantification for *Myh11-CreER*, *Pdgfra*$^{GFP:CreER}$, and *Pdgfrb*$^{CreER}$ drivers.**

**Supplemental file: R script for scRNA-seq analysis.**
