## Supplementary material for "Three-axis classification of mouse lung mesenchymal cells reveals two populations of myofibroblasts": Three-axis_scRNAseq.R

```
#R script for scRNA-seq analysis

#Load Packages
library(BiocManager)
library(Seurat)
library(dplyr)
library(Matrix)
library(ggplot2)
library(cowplot)
library(EnhancedVolcano)
library(monocle)
library(VGAM)
library(harmony)

#Cellranger aggregate was run using each individual sample output file, molecule_info.h5, from the cellranger count pipeline.
#Individual sample dataset available on: GSE180822
##https://support.10xgenomics.com/single-cell-gene-expression/software/pipelines/latest/using/aggregate

#Load Dataset and pre-process
samples <- read.csv(file.path("C:/your/file/path/", "aggregation.csv"), stringsAsFactors=F)
cells.data <- Read10X("C:/your/file/path/filtered_feature_bc_matrix/")
cells <- new("seurat", raw.data=cells.data)
cellcodes <- as.data.frame(cells@raw.data@Dimnames[[2]])
colnames(cellcodes) <- "barcodes"
rownames(cellcodes) <- cellcodes$barcodes
cellcodes$libcodes <- as.factor(gsub(pattern=".+-", replacement="", cellcodes$barcodes))
cellcodes$samples <- as.vector(samples$library_id[cellcodes$libcodes])

#Verify the head of codes
head(cellcodes)
head(samples)
sampleidentity <- cellcodes["samples"]

#Create Seurat Object
cells <- CreateSeuratObject(cells.data, meta.data=sampleidentity, min.cells=3, min.features=200, project="aggregate")
class(cells.data)
dim(cells.data)
cells.data[1:6, 1:6]
summary(colSums(cells.data))

# Check how many genes have at least one transcript in each cell
at_least_one <- apply(cells.data, 2, function(x) sum(x>0))
hist(at_least_one, breaks = 100, main = "Distribution of detected genes", xlab = "Genes with at least one tag")
hist(at_least_one, breaks = 100, main = "Distribution of detected genes",xlab = "Genes with at least one tag")
hist(colSums(cells.data), breaks = 100, main = "Expression sum per cell", xlab = "Sum expression")

#Check the number of genes detected in three or more cells
tmp <- apply(cells.data, 1, function(x) sum(x>0))
table(tmp>=3) 

#Check cells that have at least 200 detected genes
keep <- tmp>=3
tmp <- cells.data[keep,]
at_least_one <- apply(tmp, 2, function(x) sum(x>0))
summary(at_least_one)           
dim(tmp)                        
class(cells)

#Seurat analysis - Round 1
cells[["percent.mt"]] <- PercentageFeatureSet(cells, pattern = "^mt")
VlnPlot(cells, features = c("nFeature_RNA", "nCount_RNA", "percent.mt"), ncol = 3)
VlnPlot(cells, features = c("nFeature_RNA", "nCount_RNA", "percent.mt"), ncol = 3, pt.size = 0)
cells <- subset(cells, subset = nFeature_RNA > 200 & nFeature_RNA < 5000 & percent.mt < 15)
cells <- NormalizeData(cells, normalization.method = "LogNormalize", scale.factor = 10000)
cells <- FindVariableFeatures(cells, selection.method = "vst", nfeatures = 3000)
all.genes <- rownames(cells)
cells <- ScaleData(cells, vars.to.regress = "percent.mt", features = rownames(sample.data), block.size = 2000, do.par = TRUE, num.cores = 8)
cells <- RunPCA(cells, features = VariableFeatures(object = cells))
cells <- cells %>% 
  RunHarmony("samples", plot_convergence = TRUE)
cells <- RunUMAP(cells, reduction = "harmony", dims = 1:30, verbose = FALSE)
cells <- FindNeighbors(cells, reduction = "harmony", dims = 1:30, verbose = FALSE)
cells <- FindClusters(cells, verbose = FALSE) 
cells$CellType <- Idents(cells)
Idents(cells) <- "samples"
p1 <- DimPlot(cells, reduction = "umap")
Idents(cells) <- "CellType"
p2 <- DimPlot(cells, reduction = "umap", label = TRUE)
plot_grid(p1, p2)

#How many cells are in each cluster
table(Idents(cells))
write.csv(table(Idents(cells)), "C:/your/file/path/cellnumber.csv")

#How many cells are in each sample?
table(cells$samples)
write.csv(table(cells$samples), "C:/your/file/path/cellinsample.csv")

#Figure S1B Feature Plots - analysis of four cell lineages: mesenchymal, endothelial, immune and epithelial
FeaturePlot(cells, features = c("Col3a1", "Cdh5", "Ptprc", "Nkx2-1"), cols = c("lightgrey", "red"), order = TRUE)
VlnPlot(cells, features = c("Col3a1", "Cdh5", "Ptprc", "Nkx2-1"), pt.size = 0)

#Figure S1C Feature Plots - to delineate which cells express extracellular matrix genes
FeaturePlot(cells,  order = TRUE, cols = c("lightgrey", "red"), features = c("Col13a1", "Col14a1", 'Col2a1', 'Col1a1', 'Col1a2', 'Col4a1', 'Col4a2', 'Col4a3', 'Col4a4', 'Col4a5', 'Col4a6',
                                                                             'Col5a1', 'Col5a2', 'Col5a3', 'Col6a1', 'Col6a2', 'Col6a3', 'Col6a4', 'Col6a5', 'Col6a6', 'Col7a1', 'Col8a1', 'Col8a2', 
                                                                             'Col9a1', 'Col9a2', 'Col9a3', 'Col10a1', 'Col11a1', 'Col11a2', 'Col12a1', 'Col15a1', 'Col16a1', 'Col17a1', 'Col18a1', 'Col19a1', 
                                                                             'Col2a1', 'Col20a1', 'Col22a1', 'Col23a1', 'Col24a1', 'Col25a1', 'Col26a1', 'Col27a1', 'Col28a1'))

#Figure S1C Plots - to identify the location of Wnt2 fibroblasts also known as lipofibroblasts/adipofibroblasts 
FeaturePlot(cells, features = c("Plin2", "Axin2", "Wnt2", "Il33", "Meox2"), order = TRUE, cols = c("lightgrey", "red"))

#Additional anaylisis to confirm doublets and remove from object cells
doublets <- subset(cells, idents = c("25", "30", "32", "34", "35", "37"))
doublets <- NormalizeData(doublets, normalization.method = "LogNormalize", scale.factor = 10000)
doublets <- FindVariableFeatures(doublets, selection.method = "vst", nfeatures = 3000)
all.genes <- rownames(doublets)
doublets <- ScaleData(doublets, vars.to.regress = "percent.mt")
doublets <- RunPCA(doublets, features = VariableFeatures(object = doublets))
doublets <- doublets %>% 
  RunHarmony("samples", plot_convergence = TRUE)
doublets <- RunUMAP(doublets, reduction = "harmony", dims = 1:30, verbose = FALSE)
doublets <- FindNeighbors(doublets, reduction = "harmony", dims = 1:30, verbose = FALSE)
doublets <- FindClusters(doublets, verbose = FALSE, resolution = 1.6) 
doublets$CellType <- Idents(doublets)
Idents(doublets) <- "samples"
p1 <- DimPlot(doublets, reduction = "umap")
Idents(doublets) <- "CellType"
p2 <- DimPlot(doublets, reduction = "umap", label = TRUE)
plot_grid(p1, p2)

FeaturePlot(doublets, features = c("Tbx4", "Tbx5", "Col3a1", "Cdh5", "Ptprc", "Nkx2-1"))
VlnPlot(doublets, features = c("Tbx4", "Tbx5", "Col3a1", "Cdh5", "Ptprc", "Nkx2-1"), group.by = "CellType", pt.size = 0)
doublet.markers <- FindAllMarkers(doublets, only.pos = TRUE, min.pct = 0.1, logfc.threshold = 0.15) %>% group_by(cluster) %>% top_n(n = 25, wt = avg_log2FC)
write.csv(doublet.markers, "C:/your/file/path/top25_doublets.csv")
top25 <- doublet.markers %>% group_by(cluster) %>% top_n(n = 25, wt = avg_log2FC)
Idents(doublets) <- factor(Idents(doublets))
DoHeatmap(doublets, features = top25$gene, label = FALSE)

#Round2 to analyze cells that express pan-mesenchymal cell marker, Col3a1, and removed doublet clusters
Round2 <- subset(cells, idents = c("2", "3", "11", "12", "13", "18", "27", "33"))
Round2 <- NormalizeData(Round2, normalization.method = "LogNormalize", scale.factor = 10000)
Round2 <- FindVariableFeatures(Round2, selection.method = "vst", nfeatures = 3000)
Round2 <- RunPCA(Round2, features = VariableFeatures(object = Round2))
Round2 <- Round2 %>% 
  RunHarmony("samples", plot_convergence = TRUE)
Round2 <- RunUMAP(Round2, reduction = "harmony", dims = 1:30, verbose = FALSE)
Round2 <- FindNeighbors(Round2, reduction = "harmony", dims = 1:30, verbose = FALSE)
Round2 <- FindClusters(Round2, verbose = FALSE) 
Round2$CellType <- Idents(Round2)
Idents(Round2) <- "samples"
p1 <- DimPlot(Round2, reduction = "umap")
Idents(Round2) <- "CellType"
p2 <- DimPlot(Round2, reduction = "umap", label = TRUE)
plot_grid(p1, p2)

#How many cells are in each cluster?
table(Idents(Round2))
write.csv(table(Idents(Round2)), "C:/your/file/path/Round2cellnumber.csv")

#How many cells are in each sample?
table(Round2$samples)
write.csv(table(Round2$samples), "C:/your/file/path/Round2cellinsample.csv")

#Screen for mesenchymal cells and removal of undetected doublets
FeaturePlot(Round2, features = c("Tbx4", "Tbx5", "Col3a1", "Cdh5", "Ptprc", "Nkx2-1"))
VlnPlot(Round2, features = c("Tbx4", "Tbx5", "Col3a1", "Cdh5", "Ptprc", "Nkx2-1"), group.by = "CellType", pt.size = 0)

#Round3 - mesenchymal cells only
Round3 <- subset(Round2, idents = c("0", "1", "2", "3", "4", "5", "6", "7", "8", "9", "10", "11", "12", "13", "14", "15", "18"))
Round3 <- NormalizeData(Round3, normalization.method = "LogNormalize", scale.factor = 10000)
Round3 <- FindVariableFeatures(Round3, selection.method = "vst", nfeatures = 3000)
Round3 <- RunPCA(Round3, features = VariableFeatures(object = Round3))
Round3 <- Round3 %>% 
  RunHarmony("samples", plot_convergence = TRUE)
Round3 <- RunUMAP(Round3, reduction = "harmony", dims = 1:30, verbose = FALSE)
Round3 <- FindNeighbors(Round3, reduction = "harmony", dims = 1:30, verbose = FALSE)
Round3 <- FindClusters(Round3, verbose = FALSE) 
Round3$CellType <- Idents(Round3)
Idents(Round3) <- "samples"
p1 <- DimPlot(Round3, reduction = "umap")
Idents(Round3) <- "CellType"
p2 <- DimPlot(Round3, reduction = "umap", label = TRUE)
plot_grid(p1, p2)

#How many cells are in each cluster?
table(Idents(Round3))
write.csv(table(Idents(Round3)), "C:/your/file/path/Round3cellnumber.csv")

#How many cells are in each sample?
table(Round3$samples)
write.csv(table(Round3$samples), "C:/your/file/path/Round3cellinsample.csv")

#Screen for mesenchymal cells and removal of undetected doublets
FeaturePlot(Round3, features = c("Col3a1", "Cdh5", "Ptprc", "Nkx2-1"), cols = c("lightgrey", "red"), order = TRUE)
VlnPlot(Round3, features = c("Col3a1", "Cdh5", "Ptprc", "Nkx2-1"), group.by = "CellType", pt.size = 0)

#Screen known mesenchymal marker genes. 
FeaturePlot(Round3, features = c("Pdgfrb", "Pdgfra", "Fgf18", "Wnt2", "Twist2", "Acta2", "Actc1", "Msln", "Sox10","Mki67"), )
VlnPlot(Round3, features = c("Pdgfrb", "Pdgfra", "Fgf18", "Wnt2", "Twist2", "Acta2", "Actc1", "Msln", "Sox10","Mki67"), group.by = "CellType", pt.size = 0)

#Subset only mesenchymal cells - removal of immune cell clusters 18 and 20 and mesenchymal cell doublets 24 
mesenchymal <- subset(Round3, idents =  c("0", "1", "2", "3", "4", "5", "6", "7", "8", "9", "10", "11", "12", "13", "14", "15", "16", "17", "19", "21", "22", "23"))
Idents(mesenchymal) <- "samples"
samplelevels <- c("E17", "E19", "P7", "P13", "P20", "P70")
mesenchymal@meta.data$samples <- factor(x = mesenchymal@meta.data$samples, levels=c(samplelevels))
DimPlot(mesenchymal, reduction = 'umap', group.by = 'samples', cols = c('E17' = '#B79F00', 'E19' = '#00BA38', 'P7' = '#F564E3',
                                                                   'P13' = '#00BFC4', 'P20' = '#619CFF', 'P70' = '#F8766D'))
DimPlot(mesenchymal, reduction = 'umap', split.by = 'samples', cols = c('E17' = '#B79F00', 'E19' = '#00BA38', 'P7' = '#F564E3',
                                                                        'P13' = '#00BFC4', 'P20' = '#619CFF', 'P70' = '#F8766D'))

#Table S1. Analysis of top 75 differentially expressed genes per clusters and heatmap. 
Idents(mesenchymal) <- "CellType"
diff <- FindAllMarkers(mesenchymal, only.pos = TRUE, min.pct = 0.1, logfc.threshold = 0.10) %>% group_by(cluster) %>%  top_n(n = 75, wt = avg_log2FC) 
#Gene list was analyzed using http://www.informatics.jax.org/batch with additional information: Gene Ontology setting. The database was used
#to elucidate gene ontology as well as the gene details provided. 
write.csv(diff, file = "/your/path/file/TableS1.csv")

#Save DoMultiBarHeatmap function in a separate R script (found at the end of this code). Script provided by GitHub community found here: https://github.com/satijalab/seurat/issues/2201
#Acknowledgements to the user arkal who modified the code from Seurat DoHeatmap function. 
source('/your/file/path/DoMultiBarHeatmap.R')
Heatmap <- DoMultiBarHeatmap(mesenchymal, assay = "RNA", slot = "scale.data", features = diff$gene ,group.by= "CellType", additional.group.by = "samples") + theme(axis.text.y = element_text(size = 1.2))

#Re-organization of clusters based on the three-axis model proximal-distal 
my.levels <- c("4", "10", "6", "1", "13", "12", "5", "8", "9", "0", "2", "3", "7", "15", "16", "17", "23", "19", "22", "21", "11", "14")
Idents(mesenchymal) <- factor(Idents(mesenchymal), levels = c(my.levels))
DimPlot(mesenchymal, reduction = "umap", label = FALSE, cols = c('4' = '#F8766D', '10' = '#ED813E', '6' = '#DE8C00',
                                                            '1' = '#CD9600', '13' = '#B79F00', '12' = '#9DA700', '5' = '#7CAE00',
                                                            '8' = '#7CAE00', '9' = '#00BA38', '0' = '#00BE67', '2' = '#00C08B',
                                                            '3' = '#00C1A9', '7' = '#00BFC4', '15' = '#00BBDC', '16' = '#00B4F0',
                                                            '17' = '#00A9FF', '23' = '#619CFF', '19' = '#9F8CFF', '22' = '#C77CFF',
                                                            '23' = '#E36EF6', '22' = '#F564E3', '21' = '#FF61CC', '11' = '#FF64B0', 
                                                            '14' = '#FF6C91')) 

#Screen known mesenchymal marker genes. 
FeaturePlot(mesenchymal, features = c("Pdgfrb", "Pdgfra", "Fgf18", "Wnt2", "Twist2", "Acta2", "Actc1", "Msln", "Sox10","Mki67"), )
VlnPlot(mesenchymal, features = c("Pdgfrb", "Pdgfra", "Fgf18", "Wnt2", "Twist2", "Acta2", "Actc1", "Msln", "Sox10","Mki67"), group.by = "CellType", pt.size = 0)

#Find differentially expressed genes
diff2 <- FindAllMarkers(mesenchymal, only.pos = TRUE, min.pct = 0.1, logfc.threshold = 0.10) %>% group_by(cluster) %>%  top_n(n = 75, wt = avg_log2FC) 
Heatmap_axes <- DoMultiBarHeatmap(mesenchymal, assay = "RNA", slot = "scale.data", features = diff2$gene ,group.by= "CellType", additional.group.by = "samples") + theme(axis.text.y = element_text(size = 1.2))
Heatmap_axes


#Trajectory analysis without clusters proliferative clusters 11, 14, 17, 23 
trajectory <- c("4", "10", "6", "1", "13", "12", "5", "8", "9", "0", "2", "3", "7", "15", "16", "19", "22")
monocle <- subset(mesenchymal, idents = trajectory)

#Identify proliferative genes and remove them for trajectory analysis
boring.genes <- c(grep("^Gm", rownames(monocle@assays$RNA), perl=T, value=T),grep("^Ei", rownames(monocle@assays$RNA), perl=T, value=T),grep("^Dd", rownames(monocle@assays$RNA), perl=T, value=T), "Xist")
s.genes <- c("Mcm5","Pcna", "Tyms", "Fen1", "Mcm2", "Mcm4", "Rrm1", "Ung", "Gins2", "Mcm6", "Cdca7", "Dtl", "Prim1", "Uhrf1", "Mlf1ip", "Hells", "Rfc2", "Rpa2", "Nasp",  "Rad51ap1", "Gmnn", "Wdr76", "Slbp", "Ccne2", "Ubr7", "Pold3", "Msh2", "Atad2", "Rad51", "Rrm2", "Cdc45", "Cdc6", "Exo1", "Tipin", "Dscc1", "Blm", "Casp8ap2", "Usp1", "Clspn", "Pola1", "Chaf1b", "Brip1")
g2m.genes <- c("Hmgb2", "Cdk1", "Nusap1", "Ube2c", "Birc5", "Tpx2", "Top2a", "Ndc80", "Cks2", "Nuf2", "Cks1b", "Mki67", "Tmpo", "Cenpf", "Tacc3", "Fam64a", "Smc4", "Ccnb2", "Ckap2l", "Ckap2", "Aurkb", "Bub1", "Kif11", "Anp32e", "Tubb4b", "Gtse1", "Kif20b", "Hjurp", "Cdca3", "Hn1", "Cdc20", "Ttk", "Cdc25c", "Kif2c", "Rangap1", "Ncapd2", "Dlgap5", "Cdca2", "Cdca8", "Ect2", "Kif23", "Hmmr", "Aurka", "Psrc1", "Anln", "Lbr", "Ckap5", "Cenpe", "Ctcf", "Nek2", "G2e3", "Gas2l3", "Cbx5", "Cenpa")
cell.cycle <- c("Ankle1", "Cdkn3", "Cdkn1c", "Cdkn1a", "Cbx3", "Ccnb1", "Ccnd2", "Pbk", "Ccne1")

#Monocle
monocle <- NormalizeData(monocle) %>% FindVariableFeatures() %>% ScaleData()
data <- as(as.matrix(monocle@assays$RNA@data), 'sparseMatrix')
pd <- new('AnnotatedDataFrame', data=monocle@meta.data)
fData <- data.frame(gene_short_name=row.names(data),row.names = row.names(data))
fd <- new('AnnotatedDataFrame', data=fData)
HSMM <- newCellDataSet(data,
                       phenoData=pd,
                       featureData=fd,
                       expressionFamily = uninormal())
ordering_genes <- monocle@assays$RNA@var.features[1:500]
ordering_genes <- ordering_genes[!ordering_genes %in% c(boring.genes, g2m.genes, s.genes)] 
HSMM <- setOrderingFilter(HSMM, ordering_genes)
#Pull out only the genes the first 500 genes. 
HSMM <- reduceDimension(HSMM,norm_method="none", 
                        reduction_method="DDRTree",
                        max_components=2,
                        scaling=TRUE,
                        pseudo_expr=0)
HSMM <- orderCells(HSMM) #knows the difference between any two cells; arrange the time sequentially.
plot_cell_trajectory(HSMM, color_by = "samples", show_branch_points = F)
plot_cell_trajectory(HSMM, color_by = "samples", show_branch_points = F) + facet_wrap(~samples, nrow = 1)
plot_cell_trajectory(HSMM, color_by = "State", show_branch_points = T, cell_size = 1) + facet_wrap(~CellType, nrow = 1)
plot_cell_trajectory(HSMM, markers="Pdgfrb", use_color_gradient = T) + scale_color_gradient(low="lightgrey", high="red")
plot_cell_trajectory(HSMM, markers="Pdgfra", use_color_gradient = T) + scale_color_gradient(low="lightgrey", high="red")
plot_cell_trajectory(HSMM, markers="Fgf18", use_color_gradient = T) + scale_color_gradient(low="lightgrey", high="red")
plot_cell_trajectory(HSMM, markers="Acta2", use_color_gradient = T) + scale_color_gradient(low="lightgrey", high="red")
plot_cell_trajectory(HSMM, markers="Actc1", use_color_gradient = T) + scale_color_gradient(low="lightgrey", high="red")
plot_cell_trajectory(HSMM, markers="Wnt2", use_color_gradient = T) + scale_color_gradient(low="lightgrey", high="red")

#Export state (i.e. axis) monocle data into Seurat 
plot_cell_trajectory(HSMM, color_by = "State") + facet_wrap(~State, nrow = 1)
monocle$State500 <- HSMM@phenoData@data[["State"]]
Idents(monocle) <- "State500"
DimPlot(monocle, reduction = "umap", order = c('3', '1', '2'), cols = c("#CB7AFF", '#00BFC4', '#E18A00'))
DimPlot(monocle, reduction = "umap", order = c('3', '1', '2'), cols = c("#CB7AFF", '#00BFC4', '#E18A00'), split.by = 'samples')


#Vascular axis
vascular <- subset(Round4, idents = c("4", "24", "10", "21"))
vascular <- NormalizeData(vascular, normalization.method = "LogNormalize", scale.factor = 10000)
vascular <- FindVariableFeatures(vascular, selection.method = "vst", nfeatures = 3000)
vascular <- ScaleData(vascular)
vascular <- RunPCA(vascular, features = VariableFeatures(object = vascular))
vascular <- vascular %>% 
  RunHarmony("samples", plot_convergence = TRUE)
vascular <- RunUMAP(vascular, reduction = "harmony", dims = 1:30, verbose = FALSE)
vascular <- FindNeighbors(vascular, reduction = "harmony", dims = 1:30, verbose = FALSE)
vascular <- FindClusters(vascular, verbose = FALSE, resolution = 1.6) 
vascular$CellType <- Idents(vascular)
Idents(vascular) <- "samples"
p1 <- DimPlot(vascular, reduction = "umap")
Idents(vascular) <- "CellType"
p2 <- DimPlot(vascular, reduction = "umap", label = TRUE)
plot_grid(p1, p2)

#How many cells in each cluster?
table(Idents(vascular))
write.csv(table(Idents(vascular)), "/your/file/path/vascular_cellnumber.csv")

# How many cells are in each sample?
table(vascular$samples)
write.csv(table(vascular$samples), "your/file/path/vascular_cellinsample.csv")

#Find differentially expressed genes
diffvascular <- FindAllMarkers(vascular, only.pos = TRUE, min.pct = 0.1, logfc.threshold = 0.10) %>% group_by(cluster) %>%  top_n(n = 75, wt = avg_log2FC) 
write.csv(diffvascular, file = "/your/file/path/diffvascular.csv")

FeaturePlot(vascular, features = c("Pdgfrb", "Notch3", "Gap43", "Acta2", "Tagln", "Mki67"))
FeaturePlot(vascular, feature = c("Meox2", "Wnt2", "Pdgfrb", "Notch3"))

#Remove contaminating Wnt2 interstitial cells
vascular2 <- subset(vascular, idents = c("0", "1", "2", "3", "4", "5", "6", "7"))
vascular2 <- NormalizeData(vascular2, normalization.method = "LogNormalize", scale.factor = 10000)
vascular2 <- FindVariableFeatures(vascular2, selection.method = "vst", nfeatures = 3000)
vascular2 <- ScaleData(vascular2)
vascular2 <- RunPCA(vascular2, features = VariableFeatures(object = vascular2))
vascular2 <- vascular2 %>% 
  RunHarmony("samples", plot_convergence = TRUE)
vascular2 <- RunUMAP(vascular2, reduction = "harmony", dims = 1:30, verbose = FALSE)
vascular2 <- FindNeighbors(vascular2, reduction = "harmony", dims = 1:30, verbose = FALSE)
vascular2 <- FindClusters(vascular2, verbose = FALSE, resolution = 1.2) 
vascular2$CellType <- Idents(vascular2)
Idents(vascular2) <- "samples"
p1 <- DimPlot(vascular2, reduction = "umap")
Idents(vascular2) <- "CellType"
p2 <- DimPlot(vascular2, reduction = 'umap', label = FALSE, cols = c('0' = '#F8766D', '3' = '#C49A00', '1' = '#53B400',
                                                                     '5' = '#00C094', '6' = '#00B6EB', '2' = '#A58AFF', '4' = '#FB61D7'))
plot_grid(p1, p2)

#How many cells in each cluster?
table(Idents(vascular2))
write.csv(table(Idents(vascular2)), "/your/file/path/vascular2_cellnumber.csv")

# How many cells are in each sample?
table(vascular2$samples)
write.csv(table(vascular2$samples), "/your/file/path/vascular2_cellinsample.csv")

#Identify known lung mesenchymal cells associated to the vascular tree: pericytes and vascular smooth muscle. 
FeaturePlot(vascular2, features = c("Pdgfrb", "Notch3", "Gap43", "Acta2", "Tagln", "Mki67"))

#Identify differentially expressed genes based on cluster number
diffvascular2 <- FindAllMarkers(vascular2, only.pos = TRUE, min.pct = 0.1, logfc.threshold = 0.10) %>% group_by(cluster) %>%  top_n(n = 75, wt = avg_log2FC) 
write.csv(diffvascular2, file = "/your/file/path/diffvascular2.csv")

#Rename cluster numbers with cell type
vascular2@meta.data$CellType <- plyr::mapvalues(x = vascular2@meta.data$CellType,
                                                from = c("0","1","2","3","4","5","6"),
                                                to = c("Mature Pericyte", "Immature Pericyte", "Vascular Smooth Muscle", "Mature Pericyte", 
                                                       "Vascular Smooth Muscle", "Immature Pericyte", "Proliferative Pericyte"))
#Visualize UMAP with labeled clusters
Idents(vascular2) <- "CellType"
DimPlot(vascular2, reduction = 'umap', label = FALSE)

#Organize naming order for stack bar (provided by the DoMultiBarHeatmap function) and DotPlot
vascularlevels <- c("Immature Pericyte", "Mature Pericyte", "Vascular Smooth Muscle", "Proliferating Pericyte")
Idents(vascular2) <- factor(Idents(vascular2), levels = c(vascularlevels))
vascular2@meta.data$CellType <- factor(x = vascular2@meta.data$CellType, levels=c(vascularlevels))

##Table S2
diffvascular3 <- FindAllMarkers(vascular2, only.pos = TRUE, min.pct = 0.1, logfc.threshold = 0.10) %>% group_by(cluster) %>%  top_n(n = 75, wt = avg_log2FC) 
write.csv(diffvascular3, file = "/your/file/path/TableS2.csv")

#Stack bar on top of heatmap (DoMultiBarHeatmap function) 
HeatmapsampleVascular2 <- DoMultiBarHeatmap(vascular2, assay = "RNA", slot = "scale.data", features = diffvascular3$gene ,group.by= "CellType", additional.group.by = "samples") + theme(axis.text.y = element_text(size = 2.0))

#DotPlot
Plot <- rev(c("Pdgfrb", "Notch3", "Rplp0", "Rpl7a", "Gap43", "Gucy1a1", "Acta2", "Tagln", "Top2a", "Mki67"))
DotPlot(vascular3, features = Plot, cols = c("lightgrey", "red")) + coord_flip()

#Epithelial axis
epithelial <- subset(mesenchymal, idents = c("6", "1", "13", "12", "5", "8", "9", "19"))
epithelial <- NormalizeData(epithelial, normalization.method = "LogNormalize", scale.factor = 10000)
epithelial <- FindVariableFeatures(epithelial, selection.method = "vst", nfeatures = 3000)
epithelial <- ScaleData(epithelial)
epithelial <- RunPCA(epithelial, features = VariableFeatures(object = epithelial))
epithelial <- epithelial %>% 
  RunHarmony("samples", plot_convergence = TRUE)
epithelial <- RunUMAP(epithelial, reduction = "harmony", dims = 1:30, verbose = FALSE)
epithelial <- FindNeighbors(epithelial, reduction = "harmony", dims = 1:30, verbose = FALSE)
epithelial <- FindClusters(epithelial, verbose = FALSE, resolution = 8.0) 
epithelial$CellTypes <- Idents(epithelial)
Idents(epithelial) <- "samples"
p1 <- DimPlot(epithelial, reduction = "umap")
Idents(epithelial) <- "CellTypes"
p2 <- DimPlot(epithelial, reduction = "umap", label = TRUE)
plot_grid(p1, p2)

#How many cells per cluster
table(Idents(epithelial))
write.csv(table(Idents(epithelial)), "/your/file/path/epithelial2cellnumber.csv")

# How many cells are in each sample?
table(epithelial$samples)
write.csv(table(epithelial$samples), "/your/file/path/epithelial2cellinsample.csv")

cluster44 <- FindMarkers(epithelial, ident.1 = "44", only.pos = TRUE, min.pct = 0.1, logfc.threshold = 0.10)
FeaturePlot(epithelial, features = c("Myl7", "Myl3"))
write.csv(cluster44, file = "/your/file/path/cluster44.csv")
#Cluster44 are cardiac cells. 

#Analysis of lung mesenchymal cells associated to the epithelial axis. Removal of cardiac cells.
epithelial2 <- subset(epithelial, idents = c(0:43, 45))
epithelial2 <- NormalizeData(epithelial2, normalization.method = "LogNormalize", scale.factor = 10000)
epithelial2 <- FindVariableFeatures(epithelial2, selection.method = "vst", nfeatures = 3000)
epithelial2 <- ScaleData(epithelial2)
epithelial2 <- RunPCA(epithelial2, features = VariableFeatures(object = epithelial2))
epithelial2 <- epithelial2 %>% 
  RunHarmony("samples", plot_convergence = TRUE)
epithelial2 <- RunUMAP(epithelial2, reduction = "harmony", dims = 1:30, verbose = FALSE)
epithelial2 <- FindNeighbors(epithelial2, reduction = "harmony", dims = 1:30, verbose = FALSE)
epithelial2 <- FindClusters(epithelial2, verbose = FALSE, resolution = 1.4) 
epithelial2$CellTypes <- Idents(epithelial2)
Idents(epithelial2) <- "samples"
p2 <- DimPlot(epithelial2, reduction = "umap", split.by = "samples")
Idents(epithelial2) <- "CellTypes"
p1 <- DimPlot(epithelial2, reduction = "umap", label = TRUE)
plot_grid(p1, p2)

#How many cells per cluster
table(Idents(epithelial2))
write.csv(table(Idents(epithelial2)), "/your/file/path/epithelial2cellnumber.csv")

# How many cells are in each sample?
table(epithelial2$samples)
write.csv(table(epithelial2$samples), "/your/file/path/epithelial2cellinsample.csv")

#Find differentially expressed genes across clusters. 
epithelial2markers <- FindAllMarkers(epithelial2, only.pos = TRUE, min.pct = 0.1, logfc.threshold = 0.10) %>% group_by(cluster) %>% 
  top_n(n = 75, wt = avg_log2FC) 
write.csv(epithelial2markers, file = "/your/file/path/epithelial2markers.csv")

#Visualize gene expression
FeaturePlot(epithelial2, feature = c('Actc1', 'Acta2', 'Pdgfra', 'Cdh4', 'Hhip', 'Lgr6'), cols = c("lightgrey","red"), order = TRUE, split.by = "samples")

##Table S3 - Markers for each cell population of the epithelial axis
epithelial2$cell_type <- epithelial2$RNA_snn_res.1.4
epithelial2$cell_type <- plyr::mapvalues(x = epithelial2$cell_type,
                                         from = c(0,3,6,8,1,10,11,12,5,7,2,9,4,13),
                                         to = c(rep('ductal_myofibroblast',4),rep('alveolar_myofibroblast',8),rep('airway_smooth_muscle',2)))
Idents(epithelial2) <- "cell_type"
tables3 <- FindAllMarkers(epithelial2, only.pos = TRUE, min.pct = 0.1, logfc.threshold = 0.10) %>% group_by(cluster) %>% 
  top_n(n = 75, wt = avg_log2FC) 
write.csv(tables3, file = "/your/file/path/TableS3.csv")

#DotPlot
my.levels <- rev(c("Cdh4_myofibroblast", "Pdgfra_myofibroblast", "ASM"))
Plot1 <- rev(c("Hhip", "Cdh4", "Lgr6", "Tagln", "Acta2", "Myh11", "Actc1","Pdgfra")) 
Idents(epithelial2) <- factor(Idents(epithelial2), levels = c(my.levels))
DotPlot(epithelial2, features = Plot1, cols = c("grey", "red")) + coord_flip() + theme(axis.text.x = element_text(angle = 35, vjust = 0.5))


#Interstitial axis

interstitial <- subset(mesenchymal, idents = c("0", "2", "3", "7", "15", "16", "22"))
interstitial <- NormalizeData(interstitial, normalization.method = "LogNormalize", scale.factor = 10000)
interstitial <- FindVariableFeatures(interstitial, selection.method = "vst", nfeatures = 3000)
interstitial <- ScaleData(interstitial)
interstitial <- RunPCA(interstitial, features = VariableFeatures(object = interstitial))
interstitial <- interstitial %>% 
  RunHarmony("samples", plot_convergence = TRUE)
interstitial <- RunUMAP(interstitial, reduction = "harmony", dims = 1:30, verbose = FALSE)
interstitial <- FindNeighbors(interstitial, reduction = "harmony", dims = 1:30, verbose = FALSE)
interstitial <- FindClusters(interstitial, verbose = FALSE, resolution = 1.5) 
interstitial$CellType <- Idents(interstitial)
Idents(interstitial) <- "samples"
p1 <- DimPlot(interstitial, reduction = "umap", label = FALSE)
Idents(interstitial) <- "CellType"
p2 <- DimPlot(interstitial, reduction = "umap", label = TRUE)
plot_grid(p1, p2)

#How many cells per cluster
table(Idents(interstitial))
write.csv(table(Idents(interstitial)), "/your/file/path/interstitialcellnumber.csv")

# How many cells are in each sample?
table(interstitial$samples)
write.csv(table(interstitial$samples), "/your/file/path/interstitialcellinsample.csv")


#Characterize differential gene expression across 9 unlabeled clusters
interstitialmarkers <- FindAllMarkers(interstitial, only.pos = TRUE, min.pct = 0.1, logfc.threshold = 0.10) %>% group_by(cluster) %>% 
  top_n(n = 75, wt = avg_log2FC)
write.csv(interstitialmarkers, file = "/your/file/path/interstitialmarkers.csv")

#Re-analysis of interstitial cells without cluster 10 because appears to be doublets. 
#Upon comparison with clusters from epithelial axis of P13, they are very different. 
interstitial2 <- subset(interstitial, idents = c(0:9, 11))
interstitial2 <- NormalizeData(interstitial2, normalization.method = "LogNormalize", scale.factor = 10000)
interstitial2 <- FindVariableFeatures(interstitial2, selection.method = "vst", nfeatures = 3000)
interstitial2 <- ScaleData(interstitial2)
interstitial2 <- RunPCA(interstitial2, features = VariableFeatures(object = interstitial2))
interstitial2 <- interstitial2 %>% 
  RunHarmony("samples", plot_convergence = TRUE)
interstitial2 <- RunUMAP(interstitial2, reduction = "harmony", dims = 1:30, verbose = FALSE)
interstitial2 <- FindNeighbors(interstitial2, reduction = "harmony", dims = 1:30, verbose = FALSE)
interstitial2 <- FindClusters(interstitial2, verbose = FALSE, resolution = 0.8) 
interstitial2$CellType <- Idents(interstitial2)
Idents(interstitial2) <- "samples"
p1 <- DimPlot(interstitial2, reduction = "umap", label = FALSE)
Idents(interstitial2) <- "CellType"
p2 <- DimPlot(interstitial2, reduction = "umap", label = TRUE)
plot_grid(p1, p2)

#How many cells per cluster
table(Idents(interstitial2))
write.csv(table(Idents(interstitial2)), "/your/file/path/interstitial2cellnumber.csv")

# How many cells are in each sample?
table(interstitial2$samples)
write.csv(table(interstitial2$samples), "/your/file/path/interstitial2cellinsample.csv")

#Differentially expression across clusters 
interstitialmarkers2 <-FindAllMarkers(interstitial2, only.pos = TRUE, min.pct = 0.05, logfc.threshold = 0.10) %>% group_by(cluster) %>% 
  top_n(n = 400, wt = avg_log2FC) 
write.csv(interstitialmarkers2, "/your/file/path/interstitialmarkers2.csv") 

#Evaluating gene expression in featureplots
FeaturePlot(interstitial2, features = c("Meox2", "Pdgfra", "Wnt2", "Twist2", "Pi16", "Il33", "Dner", "Gli1", "Pdgfrb"),  cols = c("lightgrey", "red"), order = TRUE, )

#Cluster 9 was labeled as proximal due to its close proximity to cluster 6. Nevertheless, there might be additional 
#heterogeneity among these proximal interstitial cells. 
interstitial2$CellType <- plyr::mapvalues(x = interstitial2$CellType,
                                               from = c(0, 1, 2, 3, 4, 5, 6, 7, 8, 9), 
                                               to = c("distal","distal", "distal", "distal", "proximal", "distal", "proximal",
                                                      "distal", "distal", "proximal"))

DimPlot(interstitial2, reduction = "umap", label = FALSE)

#Table S4. Differential gene expression with assigned nomenclature.
interstitialmarkers3 <-FindAllMarkers(interstitial2, only.pos = TRUE, min.pct = 0.05, logfc.threshold = 0.10) %>% group_by(cluster) %>% 
  top_n(n = 400, wt = avg_log2FC) 
write.csv(interstitialmarkers3, "/your/file/path/TableS4.csv") 

#Comparison gene expression between proximal and distal interstitial cells. 
interstitial.levels <- c("proximal", "distal")
Plot3 <- c("Pdgfrb", "Gli1", "Dner", "Il33", "Pi16", "Twist2", "Wnt2", "Pdgfra", "Meox2")
Idents(interstitial2) <- factor(Idents(interstitial2), levels = rev(c(interstitial.levels)))
DotPlot(interstitial2, features = rev(Plot3), cols = c("lightgrey", "red"),) + theme(axis.text.x = element_text(angle = 45, hjust = 1))

#Find the difference between young and adult Wnt2 population
interstitial2$cluster.samples <- paste(interstitial2$CellType, interstitial2$samples, sep = "_")
interstitial2@meta.data
interstitial2$Wnt2 <- plyr::mapvalues(x = interstitial2$CellType,
                                      from = c(0, 1, 2, 3, 4, 5, 6, 7, 8, 9), 
                                      to = c("young","adult", "young", "young", "proximal", "adult", "proximal",
                                             "young", "young", "proximal"))
Idents(interstitial2) <- "Wnt2"
cluster <- FindMarkers(interstitial2, ident.1 = "adult", ident.2 = "young")
EnhancedVolcano(cluster, lab = rownames(cluster), x='avg_log2FC', y='p_val_adj', FCcutoff = 0.9, pCutoff = 10e-4, 
                xlim=c(-max(abs(cluster$avg_log2FC))-.1,max(abs(cluster$avg_log2FC))+.1), drawConnectors = TRUE,
                gridlines.major = F, gridlines.minor = F, xlab = "ln(fold change)", colAlpha = 1,
                title=NULL, subtitle=NULL, col=c('black','black','black','red'))


##DoMultiBarHeatmap function from github: https://github.com/satijalab/seurat/issues/2201 by arkal user
##Create a new R script strictly for the code and source it to R environment. 

suppressPackageStartupMessages({
  library(rlang)
})

DoMultiBarHeatmap <- function (object, 
                               features = NULL, 
                               cells = NULL, 
                               group.by = "ident", 
                               additional.group.by = NULL, 
                               additional.group.sort.by = NULL, 
                               cols.use = NULL,
                               group.bar = TRUE, 
                               disp.min = -2.5, 
                               disp.max = NULL, 
                               slot = "scale.data", 
                               assay = NULL, 
                               label = TRUE, 
                               size = 5.5, 
                               hjust = 0, 
                               angle = 45, 
                               raster = TRUE, 
                               draw.lines = TRUE, 
                               lines.width = NULL, 
                               group.bar.height = 0.02, 
                               combine = TRUE) 
{
  cells <- cells %||% colnames(x = object)
  if (is.numeric(x = cells)) {
    cells <- colnames(x = object)[cells]
  }
  assay <- assay %||% DefaultAssay(object = object)
  DefaultAssay(object = object) <- assay
  features <- features %||% VariableFeatures(object = object)
  ## Why reverse???
  features <- rev(x = unique(x = features))
  disp.max <- disp.max %||% ifelse(test = slot == "scale.data", 
                                   yes = 2.5, no = 6)
  possible.features <- rownames(x = GetAssayData(object = object, 
                                                 slot = slot))
  if (any(!features %in% possible.features)) {
    bad.features <- features[!features %in% possible.features]
    features <- features[features %in% possible.features]
    if (length(x = features) == 0) {
      stop("No requested features found in the ", slot, 
           " slot for the ", assay, " assay.")
    }
    warning("The following features were omitted as they were not found in the ", 
            slot, " slot for the ", assay, " assay: ", paste(bad.features, 
                                                             collapse = ", "))
  }
  
  if (!is.null(additional.group.sort.by)) {
    if (any(!additional.group.sort.by %in% additional.group.by)) {
      bad.sorts <- additional.group.sort.by[!additional.group.sort.by %in% additional.group.by]
      additional.group.sort.by <- additional.group.sort.by[additional.group.sort.by %in% additional.group.by]
      if (length(x = bad.sorts) > 0) {
        warning("The following additional sorts were omitted as they were not a subset of additional.group.by : ", 
                paste(bad.sorts, collapse = ", "))
      }
    }
  }
  
  data <- as.data.frame(x = as.matrix(x = t(x = GetAssayData(object = object, 
                                                             slot = slot)[features, cells, drop = FALSE])))
  
  object <- suppressMessages(expr = StashIdent(object = object, 
                                               save.name = "ident"))
  group.by <- group.by %||% "ident"
  groups.use <- object[[c(group.by, additional.group.by[!additional.group.by %in% group.by])]][cells, , drop = FALSE]
  plots <- list()
  for (i in group.by) {
    data.group <- data
    if (!is_null(additional.group.by)) {
      additional.group.use <- additional.group.by[additional.group.by!=i]  
      if (!is_null(additional.group.sort.by)){
        additional.sort.use = additional.group.sort.by[additional.group.sort.by != i]  
      } else {
        additional.sort.use = NULL
      }
    } else {
      additional.group.use = NULL
      additional.sort.use = NULL
    }
    
    group.use <- groups.use[, c(i, additional.group.use), drop = FALSE]
    
    for(colname in colnames(group.use)){
      if (!is.factor(x = group.use[[colname]])) {
        group.use[[colname]] <- factor(x = group.use[[colname]])
      }  
    }
    
    if (draw.lines) {
      lines.width <- lines.width %||% ceiling(x = nrow(x = data.group) * 
                                                0.0025)
      placeholder.cells <- sapply(X = 1:(length(x = levels(x = group.use[[i]])) * 
                                           lines.width), FUN = function(x) {
                                             return(Seurat:::RandomName(length = 20))
                                           })
      placeholder.groups <- data.frame(rep(x = levels(x = group.use[[i]]), times = lines.width))
      group.levels <- list()
      group.levels[[i]] = levels(x = group.use[[i]])
      for (j in additional.group.use) {
        group.levels[[j]] <- levels(x = group.use[[j]])
        placeholder.groups[[j]] = NA
      }
      
      colnames(placeholder.groups) <- colnames(group.use)
      rownames(placeholder.groups) <- placeholder.cells
      
      group.use <- sapply(group.use, as.vector)
      rownames(x = group.use) <- cells
      
      group.use <- rbind(group.use, placeholder.groups)
      
      for (j in names(group.levels)) {
        group.use[[j]] <- factor(x = group.use[[j]], levels = group.levels[[j]])
      }
      
      na.data.group <- matrix(data = NA, nrow = length(x = placeholder.cells), 
                              ncol = ncol(x = data.group), dimnames = list(placeholder.cells, 
                                                                           colnames(x = data.group)))
      data.group <- rbind(data.group, na.data.group)
    }
    
    order_expr <- paste0('order(', paste(c(i, additional.sort.use), collapse=','), ')')
    group.use = with(group.use, group.use[eval(parse(text=order_expr)), , drop=F])
    
    plot <- Seurat:::SingleRasterMap(data = data.group, raster = raster, 
                                     disp.min = disp.min, disp.max = disp.max, feature.order = features, 
                                     cell.order = rownames(x = group.use), group.by = group.use[[i]])
    
    if (group.bar) {
      pbuild <- ggplot_build(plot = plot)
      group.use2 <- group.use
      cols <- list()
      na.group <- Seurat:::RandomName(length = 20)
      for (colname in rev(x = colnames(group.use2))) {
        if (colname == i) {
          colid = paste0('Identity (', colname, ')')
        } else {
          colid = colname
        }
        
        # Default
        cols[[colname]] <- c(scales::hue_pal()(length(x = levels(x = group.use[[colname]]))))  
        
        #Overwrite if better value is provided
        if (!is_null(cols.use[[colname]])) {
          req_length = length(x = levels(group.use))
          if (length(cols.use[[colname]]) < req_length){
            warning("Cannot use provided colors for ", colname, " since there aren't enough colors.")
          } else {
            if (!is_null(names(cols.use[[colname]]))) {
              if (all(levels(group.use[[colname]]) %in% names(cols.use[[colname]]))) {
                cols[[colname]] <- as.vector(cols.use[[colname]][levels(group.use[[colname]])])
              } else {
                warning("Cannot use provided colors for ", colname, " since all levels (", paste(levels(group.use[[colname]]), collapse=","), ") are not represented.")
              }
            } else {
              cols[[colname]] <- as.vector(cols.use[[colname]])[c(1:length(x = levels(x = group.use[[colname]])))]
            }
          }
        }
        
        # Add white if there's lines
        if (draw.lines) {
          levels(x = group.use2[[colname]]) <- c(levels(x = group.use2[[colname]]), na.group)  
          group.use2[placeholder.cells, colname] <- na.group
          cols[[colname]] <- c(cols[[colname]], "#FFFFFF")
        }
        names(x = cols[[colname]]) <- levels(x = group.use2[[colname]])
        
        y.range <- diff(x = pbuild$layout$panel_params[[1]]$y.range)
        y.pos <- max(pbuild$layout$panel_params[[1]]$y.range) + y.range * 0.015
        y.max <- y.pos + group.bar.height * y.range
        pbuild$layout$panel_params[[1]]$y.range <- c(pbuild$layout$panel_params[[1]]$y.range[1], y.max)
        
        plot <- suppressMessages(plot + 
                                   annotation_raster(raster = t(x = cols[[colname]][group.use2[[colname]]]),  xmin = -Inf, xmax = Inf, ymin = y.pos, ymax = y.max) + 
                                   annotation_custom(grob = grid::textGrob(label = colid, hjust = 0, gp = gpar(cex = 0.75)), ymin = mean(c(y.pos, y.max)), ymax = mean(c(y.pos, y.max)), xmin = Inf, xmax = Inf) +
                                   coord_cartesian(ylim = c(0, y.max), clip = "off")) 
        
        if ((colname == i) && label) {
          x.max <- max(pbuild$layout$panel_params[[1]]$x.range)
          x.divs <- pbuild$layout$panel_params[[1]]$x.major %||% pbuild$layout$panel_params[[1]]$x$break_positions()
          group.use$x <- x.divs
          label.x.pos <- tapply(X = group.use$x, INDEX = group.use[[colname]],
                                FUN = median) * x.max
          label.x.pos <- data.frame(group = names(x = label.x.pos), 
                                    label.x.pos)
          plot <- plot + geom_text(stat = "identity", 
                                   data = label.x.pos, aes_string(label = "group", 
                                                                  x = "label.x.pos"), y = y.max + y.max * 
                                     0.03 * 0.5, angle = angle, hjust = hjust, 
                                   size = size)
          plot <- suppressMessages(plot + coord_cartesian(ylim = c(0, 
                                                                   y.max + y.max * 0.002 * max(nchar(x = levels(x = group.use[[colname]]))) * 
                                                                     size), clip = "off"))
        }
      }
    }
    plot <- plot + theme(line = element_blank())
    plots[[i]] <- plot
  }
  p1 <- plot + theme(line = element_blank())
  p2 <- plots[[i]]
  
  p1 + p2
  
}
return(p1 + p2)
```
